# Ara h 2 and FcεRIα binding elicit state-dependent allostery in IgE

**DOI:** 10.64898/2026.09.11.750994

**Authors:** Mathialagan Mugundhan, Qinyu Jia, Palur V Raghuvamsi, Su Ning Loh, Alicia Ghia Min Ong, Krithika Subramani, Kamolrat Somboon, Firdaus Samsudin, Zhong-Ping Yao, Radoslaw Mikolaj Sobota, Samuel Ken-En Gan, Peter J Bond, Zheng Ser, Chinh Tran-To Su

## Abstract

**Background:** Peanuts are among the most prevalent food allergens responsible for anaphylaxis, particularly in children. Allergic responses may be mitigated by disrupting molecular interactions between immunoglobulin-E (IgE) and its binding partners: peanut allergen (Ara h 2) and IgE receptors (FcεRIa). However, the structural dynamics of IgE upon engaging these factors are not yet fully elucidated.

**Objective:** To characterize how Ara h 2 and/or FcεRIα binding influence IgE flexibility and its interdomain communications.

**Methods:** The structural dynamics of full-length IgE in various unbound and bound states were examined by combining molecular dynamics (MD) simulations with cross-linking mass spectrometry (XLMS). Causal relationship between IgE domains were characterized to infer allosteric communication pathways, which were assessed through hydrogen-deuterium exchange mass spectrometry (HDX-MS).

**Results:** Non-canonical bent IgE conformations were identified. We demonstrate that FcεRIα binding immobilizes IgE predominantly by stabilizing and restricting Fc flexibility, whereas Ara h 2 binding induces specific conformational changes within the FcεRIα-binding region. Our results further suggest that the Cε domains retain sufficient freedom to enable causal dynamics by the Fabs in a binding-dependent manner, thereby modulating the receptor engagement potential of IgE. Notably, concurrent binding of Ara h 2 and FcεRIα to IgE shifts the allosteric communication between the Fabs and Fc regions from the Cε3 to Cε4 domain.

**Conclusion:** Our findings provide new insights into the dynamic regulation of IgE and its role in allergic responses. This could serve as a structural framework for understanding IgE signaling in peanut allergy and suggest potential targets for therapeutic interventions.

**Key messages:**

- IgE adopts non-canonical bent FcεRIα-bound conformations with unexpected interactions between Fabs and FcεRIα
- More diverse IgE-Ara h 2 interactions were observed in the presence of FcεRIα, indicating receptor-mediated stabilization of alternative binding modes
- Binding to Ara h 2 promotes a more compact Fc structure resembling the FcεRIα-bound state, suggesting antigen binding primes IgE for receptor interaction.
- Allosteric signaling shifts depending on binding state: in Ara h 2-bound IgE, signaling is mediated through Cε3 and Cε4, whereas in FcεRIα-bound states, signaling shifts toward Cε4. Concurrent Ara h 2 and FcεRIα binding results in a “hub-dominated” network with fewer but stronger interdomain dependencies, integrating inputs from Fab movements.

**Capsule summary:** This study provides molecular insight into the cooperative mechanisms within IgE underlying allergic responses. The binding-dependent allostery highlights potential targets for reducing receptor activation and may inform development of more precise anti-IgE therapies for managing allergies.

## INTRODUCTION

Several food proteins, including peanut allergens, can trigger severe immune reactions^1–3^. Unlike many food allergies that resolve with age, peanut allergy often persists throughout life and remains common especially in Western countries, where its prevalence has been increasing^4^. Among peanut allergens, Ara h 2 is the most potent and accounts for allergenicity in 80-90% of affected individuals^5,6^. These allergic responses are mediated by immunoglobulin-E (IgE) through its interactions with allergens such as Ara h 2 and the immune receptor FcεR^7,8^.

IgE is the primary antibody isotype associated with Type-I hypersensitivity and mast cell or basophil activation^9,10^. Cellular activation results from multivalent allergen binding^9^ to IgE attached to the high-affinity extracellular domain of the FcεRI α-subunit (FcεRIα). Disrupting these interactions may provide a strategy for modulating allergic responses, potentially managing peanut allergy^11^. Our previous study^12^ suggested that prolonged FcεRIα engagement promotes persistent IgE states with a heterogeneous microenvironment containing more dissociative IgE, likely reflecting contributions from multiple IgE regions^12^.

Despite immunological advances, structural studies of IgE remain limited, especially for its interactions with antigen and receptor that govern allergic responses. Determining high-resolution structures of full-length IgE complexed with allergens and/or receptors remains challenging, due to the large assembly size, Fab-Fc interdomain flexibility^13^, and heterogeneous glycosylation of both IgE and FcεR^14,15^. Therefore, future allergen-specific IgE studies and therapeutic IgE engineering would benefit from a holistic view of intact IgE rather than isolated domains alone^16^, alongside its antigen and receptor.

IgE consists of paired heavy and light chains with variable and constant domains (VH and Cε1–Cε4 on the heavy; VL and Cκ/Cλ on the light). Its Cε3 domains bind FcεRIα^17,18^, while Fab regions recognize allergens such as Ara h 2. Existing structural data have largely been derived from isolated IgE domains rather than intact IgE-allergen-FcεRIα complexes. These studies^18,19^ showed that IgE adopts bent or extended conformations, becoming more acutely bent when bound to FcεRIα, but can become partially or fully extended in the presence of anti-ε-chain Fab such as omalizumab^20^ or aεFab^21^, demonstrating the remarkable conformational plasticity of IgE. However, since antigen binding was not examined, it remains unresolved how antigen engagement affects the conformational changes of FcεRIα-bound IgE and how domain transitions propagate to FcεRIα engagement.

To address this, we examined full-length IgE dynamics in unbound and FcεRIα-bound states in the presence of Ara h 2 by combining molecular dynamics (MD) simulations and cross-linking mass spectrometry (XLMS), which has been useful for characterizing antibody-antigen interactions^22–24^. We also analyzed causal relationships between IgE domains and assessed inferred allosteric pathways with hydrogen-deuterium exchange mass spectrometry (HDX-MS), providing new insights into how Ara h 2 and FcεRIα binding shape IgE structural flexibility.

## METHODS

### Modeling unbound bent/extended IgE and FcεRIα-bound bent IgE structures

Templates to model bent and extended IgE were referenced from IgE-Fc scaffolds^12^ (Cε1-Cε4) and from PDB:6EYO, respectively. Cκ was used as light chain constant domain. Fv sequences were substituted with fragments derived from published sequences (PA12C07) known for high allergenicity to peanut allergens, identified by Croote et. al^25^. These Fv models, particularly CDR-H3, were generated using ROSIE^26^. Full-length IgE models were reconstituted using MODELLER v10.1^27^. The FcεRIα-bound bent IgE complex was generated by superimposing FcεRIα at the FcεRIα-binding site located at the Cε3-Cε3 domains using PDB:2Y7Q as a template. Glycans were modelled using CHARMM-GUI^28^. These glycosylated models underwent MD simulations (500 ns × 3 replicates/each). See Supplementary File for full details.

### Modeling Ara h 2-bound IgE complexes

Ara h 2 was extracted from PDB:3OB4 (residues 1028-1148). Missing residues were added using MODELLER v10.1^27^ and renumbered according to ref^6^. Region-I (^22^RRCQSQLERANLRPCE^37^) and region-II (^34^RPCEQHLMQKIQR^46^) were referenced from previous studies^6,29^. Interacting residues were determined by WHISCY^30^ and used as “active residues” to dock to IgE-Fv models using HADDOCK v2.4 webserver^31^ with default settings. HDX-MS showed Ara h 2 interactions involved the ^113^QQIMENQSDRLQGRQQEQQ^1^^31^ fragment, and hence docked Ara h 2-IgE^Fv^ complexes compatible with the HDX-MS data were used as starting points for further simulations, applying similar protocols as described above.

### Structural Equation Model (SEM) with Directed Acyclic Graph (DAG) for causal inference analysis

Distances of each Fab to Cε domains, local Cε2/Cε3/Cε4, Cε2-Cε3 linkers (P338, R339, and G340)^17,21^ and the FcεRIα-binding site, specifically involving direct contacts R339/D367 on chain A and P431 on chain C (referenced from PDB:2Y7Q^18^ and 1F6A^17^) were used to define a DAG for each IgE states. Causal relationships were learned using an ensemble algorithm of constraint-based PC (Peter-Clark) from causal-learn^32^, LiNGAM^33^, and NoTears from causalnex^34^; each was executed across 300 bootstrap resamples to assess stability of inferred causality. The resulting edges were subsequently aggregated to identify a consensus DAG topology (edge_threshold=0.6), retaining only edges consistently recovered across the algorithms and the resamples to avoid algorithm-specific biases. Non-linear SEM was applied to fit onto the constructed DAGs. *RandomForestRegressor* (n_estimators=200, max_depth=6) was used.

### Experimental preparation of IgE, Ara h 2, and FcεRIα

IgE was produced and purified (by GenScript USA Inc., with Lot Number: U138NHK250-4/P9IB001) using published VH/VL sequences (PA12C07)^25^. Constant domains included light chain Cκ and heavy chain Cε of human IgE. Peanut allergen Ara h 2 was purchased from LifeSpan BioSciences (Cat ID: LS-G21762) and FcεRIα from AcroBioSystems (Cat No: FCA-H5228). Binding abilities of Ara h 2 to IgE, of FcεRIα to IgE, and FcεRIα-IgE-Ara h 2 were confirmed by GenScript USA Inc. using the Octet® BLI Discovery version 12.2.2.26. See Supplementary File for full details.

### Cross-linking Mass Spectrometry

Cross-linking with 2mM DSSO was performed for 4 different mixtures (IgE: Arah2 1:1, IgE: FcεRIα 1:1, IgE: Arah2: FcεRIα 1:2:1, IgE: Arah2: FcεRIα 2:1:2) and confirmed by Colloidal Blue staining of SDS-PAGE gel. Cross-linked proteins were reduced with 10 mM Tris(2-carboxyethyl) phosphine (TCEP), alkylated with 55 mM chloroacetamide (CAA) and digested with LysC (1:25 w/w) for 4h at 25°C followed by GluC (1:25, w/w) overnight at 25°C. Digested peptides were desalted using C18 StageTips and vacuum centrifuged. Dried peptides were resuspended and analyzed on Orbitrap Lumos mass spectrometer coupled to an EASY-nLC 1200 system. MS1 acquisition was performed in Orbitrap with MS2 CID 30% - MS2 HCD 35% fragmentation. Cross-linked peptide search was performed using MeroX software^35^ (version 2.0.1.4) with a false discovery rate (FDR) of 1% against database of Arah2, IgE heavy and light chains, and FcεRIα sequences. See Supplementary File for full details.

### Hydrogen-Deuterium Exchange Mass Spectrometry (HDX-MS)

HDX-MS experiments were performed in four setups: apo Ara h 2 (90pmol), apo IgE (90pmol), Ara h 2-IgE (180pmol: 90pmol), and FcεRIα-IgE-Ara h 2 (90pmol: 90pmol:180pmol). For complex formation, Ara h 2 and IgE or FcεRIα-IgE-Ara h 2 were pre-incubated at room temperature for 30 min prior to deuterium labeling. All labeling (1, 10, and 100 min) experiments were performed in triplicate, and reported deuterium uptake values represent the mean of replicates without correction for back-exchanges. See Supplementary File for full details.

## RESULTS

### Observed structural differences across various conformational states of free IgE

Full-length IgE dynamics were simulated in unbound extended (*Uextd*), unbound bent (*Ubent*), and FcεRIα-bound bent (*FcR-bent*) states to examine domain motions associated with FcεRIα engagement (Figure 1). The extended conformation was excluded from FcεRIα-bound simulations because its FcεRIα-binding site is concealed within the Cε2-Cε3/4 region, preventing interaction^17,36^.

**Figure 1:**
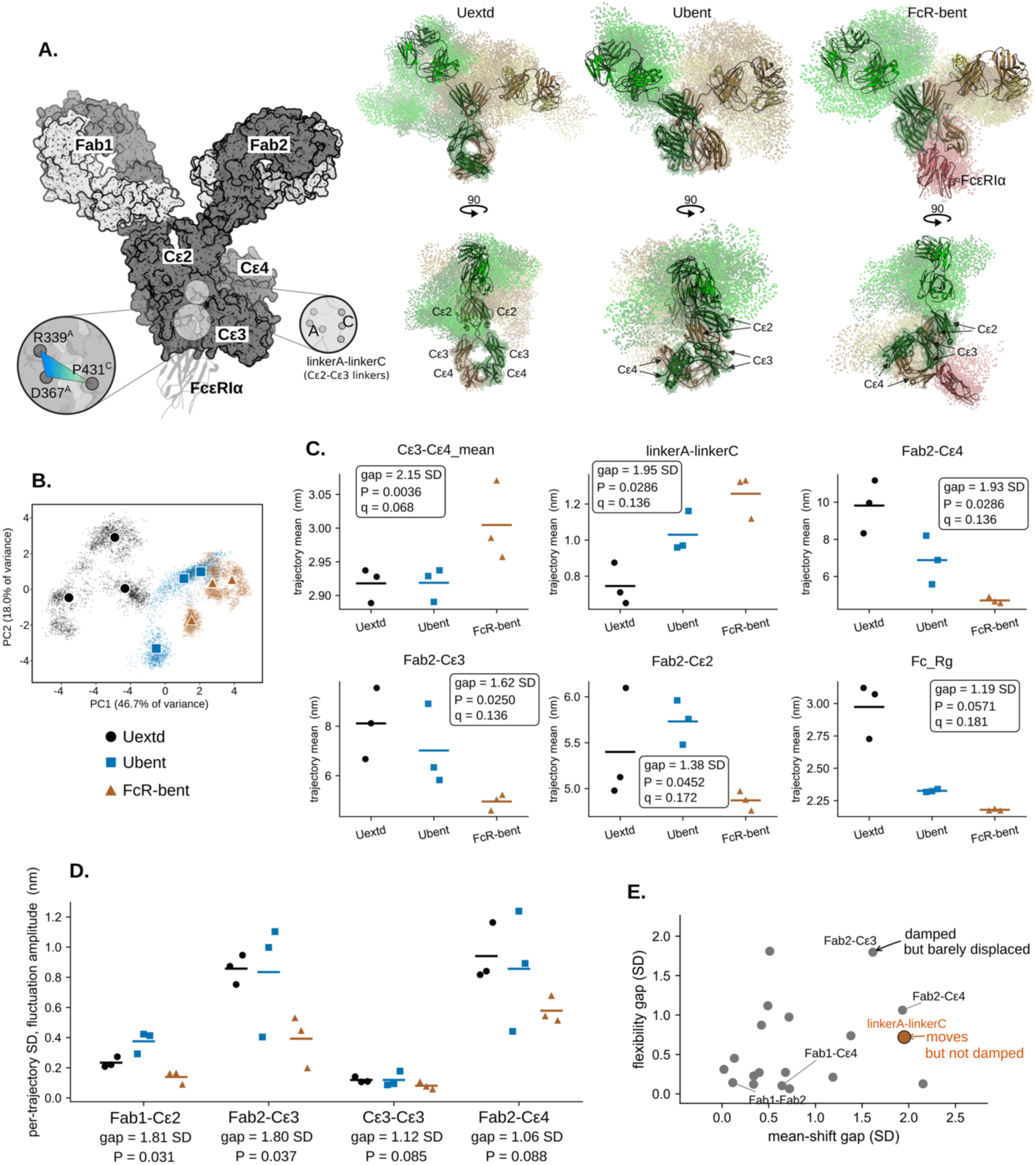
Observed structural differences across various conformational states of free IgE: unbound extended *Uextd*, unbound bent *Ubent*, and FcεRIα-bound bent *FcR-bent*. **(A)** Illustration of IgE structure with Fab and constant domains. The FcεRIα-binding region involving R339, D367 (chain A), and P431 (chain C) and the inter-chain Cε2-Cε3 linkers (*linkerA-linkerC*) are highlighted. Structural representatives (with equilibrated models in bold) are shown to illustrate asymmetrical Fabs movements across the IgE states. **(B)** Principal component analysis projecting every frame from the three IgE states to capture the most structural variance. **(C)** Six descriptors that display the most differences (*gap*, unit in SD: standard deviation) between the *FcR-bent* and both the unbound states. Each state contains n=3 trajectories, *P* and *q* are p-values of the permutation test and after the Benjamini-Hochberg false discovery rate (FDR) correction for testing 19 observables.

Nineteen structural descriptors, including domain distances and angles, were analyzed across these IgE states. The *FcR-bent* was structurally distinct from both the unbound ensembles (PC1 = 46.7%, PC2 = 18.0% variance; Figure 1B). The two unbound ensembles were more strongly separated, likely reflecting differences in their starting structures, including Q376N in *Uextd*. Notably, *FcR-bent* occupied conformational space resembled an extension of a potential *Uextd*-to-*Ubent* transition. Despite capturing ∼65% of total variance, this ordination suggests FcεRIα engagement shifts the Fc conformational ensemble. FcεRIα engagement was associated with increased Cε3-Cε4 separation and widening of the inter-chain Cε2-Cε3 linkers (linkerA-linkerC), coupled with compaction of the Fab2 arm toward the Fc core (Figure 1C). These changes indicated localized repositioning of the Cε3-proximal hinge region rather than a rigid-body rearrangement of the whole Fc. Compared to both unbound states, the *FcR-bent* ensemble exhibited larger Cε3-Cε4 (gap = 2.15 SD, P = 0.0036) and linkerA-linkerC distances (gap = 1.95 SD, P = 0.029), reduced Fab2-Cε distances, and smaller Fc radius of gyration. The FcεRIα engagement also restrained Fab mobility independently of its effect on Fc architecture (Figure 1D). Smaller per-trajectory fluctuations in Fab1-Cε2 and Fab2-Cε3 in the *FcR-bent* indicated constrained Fab-arm excursions. Comparison of mean positional shifts and flexibility amplitudes (Figure 1E) showed these effects to be largely independent, e.g. Fab2-Cε3 displayed substantial damping with minimal displacement whereas linkerA-linkerC flexibility showed an opposite pattern. Although none of the 19 descriptors individually remained significant after Benjamini-Hochberg correction (Figure S1), the broad and directionally coherent changes across multiple Fc and Fab2 proximities supports a mechanistically plausible FcεRIα-associated shift in the Fc/Fab conformational ensemble. Nevertheless, these findings should be considered as structural hypothesis rather than definitive evidence due to limited sampling.

### Non-canonical conformations of IgE when engaging Ara h 2 and/or FcεRIα

Multiple linear^6,37,38^ and conformational^6,39^ epitopes have been identified on Ara h 2 (Figure 2A). To examine diverse IgE recognition, three antigenic regions were examined. Region-I overlaps known linear (^22^RRCQSQLER^31^) and conformational (^25^QS--ERANLRP-E^37^) epitopes, whereas region-II contains a linear epitope (^34^RPCEQHLMQ^42^); both form adjacent IgE-binding interfaces^29,39^. These Ara h 2-bound IgE complexes were generated by docking IgE^Fab^^1^ models to each interface and evaluated using MD simulations.

**Figure 2:**
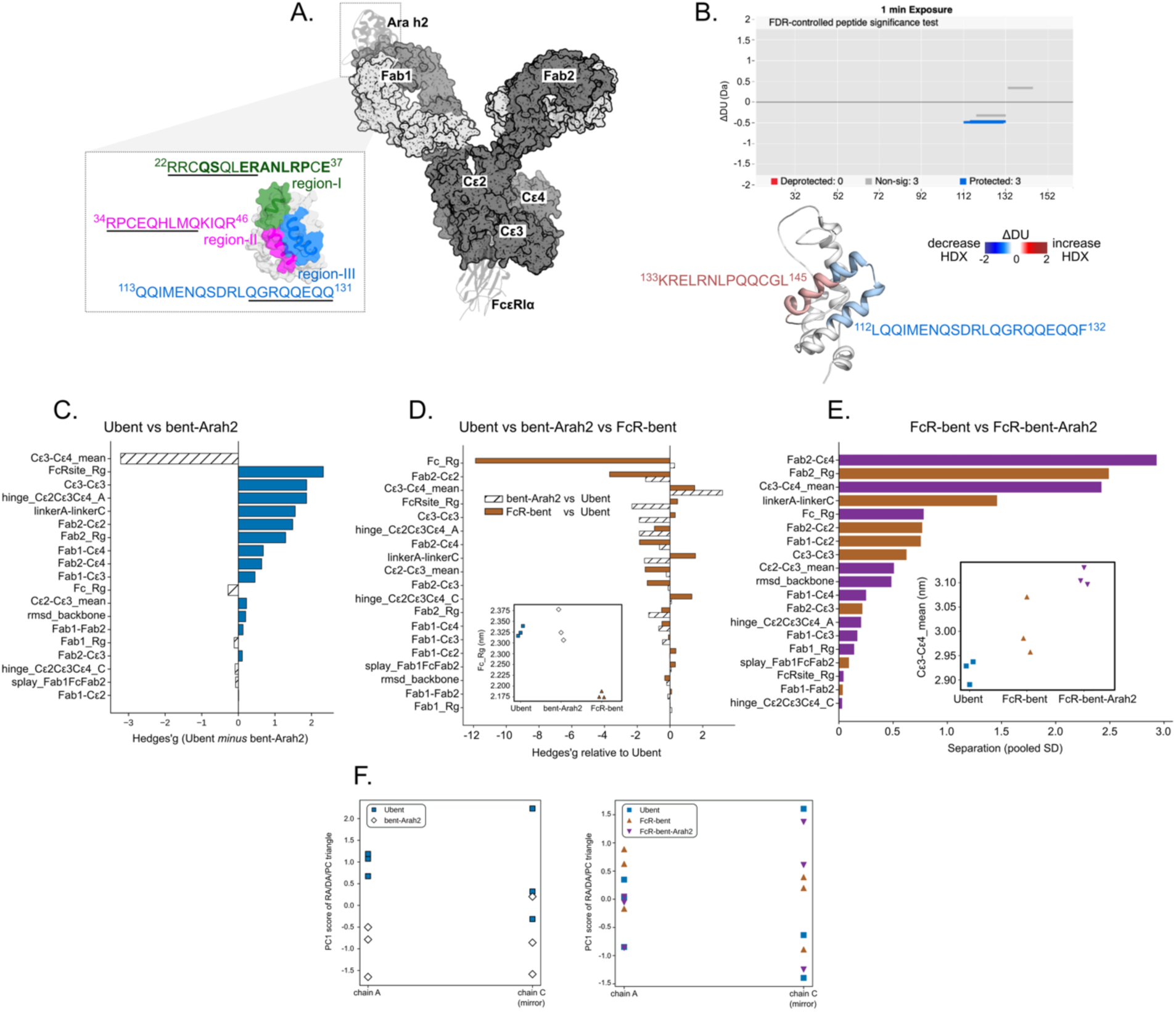
Conformational changes of IgE in the presence of the peanut allergen Ara h 2 and/or FcεRIα. **(A)** Illustration of Ara h 2-FcεRIα bound IgE structure with various Fab and constant domains. The Ara h 2 structure encompassing the three antigenic regions (with known epitopes underlined) is shown. **(B)** A Woods plot shows differences in deuterium exchange (ΔDU) within Ara h 2 calculated from the IgE-bound state relative to the free Ara h 2 state at 1-minute deuterium labelling time. The length of each line represents the corresponding pepsin proteolyzed Ara h 2 peptides, shown along x-axis from N- to C-termini. A 99% confidence interval (CI) was used to analyze each labelling time. Peptides showing differences greater than CI are highlighted as per color key. Corresponding differences in deuterium uptake were mapped on to the Ara h 2 structure for the 1 min labelling time. **(C-E)** Statistical analyses of 19 structural descriptors compared across unbound bent (*Ubent*), Ara h 2-bound (*bent-Arah2),* FcεRIα-bound (*FcR-bent*), and the concurrent bound (*FcR-bent-Arah2*) IgE states. In each comparison, the state with higher value in each measurement is colored accordingly. The Hedge’s g (standardized effect size) is used to express the gap between two group means. **(F)** PC1 scores per trajectory (n=3) of the FcεRIα-binding site triangle (R^A^/D^A^/P^C^) are plotted for chain A and chain C (mirrored) to demonstrate the asymmetrical binding properties of the FcεRIα-binding site.

Region-I showed intermittent binding, whereas region-II bound consistently and formed more contacts with IgE^Fab1^ (Table S1, Figure S2). Simultaneous engagement of both regions was not observed (Figure S3). Overall, IgE^Fab1^ binding increased Ara h 2 flexibility in unstructured regions while preserving stable helical structure (Figure S4-S5).

Several IgE-Ara h 2 complexes also involved region-III (including epitope ^124^QGRQQEQQ^1^^31^), as supported by HDX-MS data (Figure 2B). Decreased deuterium exchange across residues 113-132 upon IgE binding confirmed the engagement at region-III. Additionally, residues 133–145 (*red* in Figure 2B and S6) showed increased exchange, indicating that IgE binding induces Ara h 2 local structural rearrangements and increased solvent exposure adjacent to the epitope.

To further assess such effects on IgE, simulations were performed for region-III-bound Ara h 2 to full-length free IgE (*bent-Arah2*) and to FcεRIα-bound IgE (*FcR-bent-Arah2*). Ara h 2 binding was associated with coordinated Fc rearrangements near the FcεRIα-binding region (Figure 2C). Compared to *Ubent*, *bent-Arah2* showed increased Cε3-Cε4 separation (∼2.92 to ∼3.01 nm) and reduced Cε2-Cε3-Cε4 angle on the Ara h 2-engaged chain A (112.3° to 107.1°). Meanwhile, the FcRsite_Rg, linkerA-linkerC, and the Cε3-Cε3 distance decreased while Fab2 moved closer to Cε2 (∼5.73 to ∼5.17 nm) and became more compact (Fab2_Rg: ∼2.21 to ∼1.92 nm). Because *Ubent* and *bent-Arah2* were constructed from the same IgE template, these structural shifts were mainly attributed to the Ara h 2 engagement.

Comparison with *FcR-bent* indicated that the Ara h 2-induced remodeling appears structurally distinct from the FcεRIα-induced changes rather than an attenuated structural mimic (Figure 2D). The FcεRIα binding exhibited a pronounced global Fc compactness. Several observables including Fc_Rg, FcRsite_Rg, linkerA-linkerC, and Cε3-Cε3 distance shifted in opposite directions under Ara h 2 vs FcεRIα binding –with Ara h 2 producing larger perturbation.

Concurrent FcεRIα and Ara h 2 engagement yielded non-additive effects. Although Fab2-Cε4 distance decreased in *FcR-bent* relative to *Ubent* (∼4.71 vs ∼6.9 nm), subsequent Ara h 2 binding increased this distance to ∼7.02 nm, suggesting a partial reversal. The R^A^/D^A^/P^C^ contact triangle showed a similar non-monotonic response, reaching its most expanded state in *FcR-bent* and partially reducing toward the baseline as in *Ubent* upon Ara h 2 co-engagement (Figure 2F). In contrast, Cε3-Cε4 distance increased monotonically across *Ubent*, *FcR-bent*, and *FcR-bent-Arah2*, indicating a cumulative effect at this interface (Figure 2E). Together, these observations suggest that Ara h 2-mediated Fc remodeling is dependent on FcεRIα occupancy and conformational state of IgE.

Cross-linking mass spectrometry (XLMS) was performed to map IgE interactions across four complexes: IgE-Ara h 2, IgE-FcεRIα, IgE-FcεRIα-[Ara h 2]_2_ (1:1:2), and [IgE]_2_-[FcεRIα]_2_-Ara h 2 (2:2:1). Cross-linking was confirmed by SDS-PAGE (Figure 3A), showing higher molecular-weight species (≥180 kDa) upon DSSO^40^ treatment.

**Figure 3:**
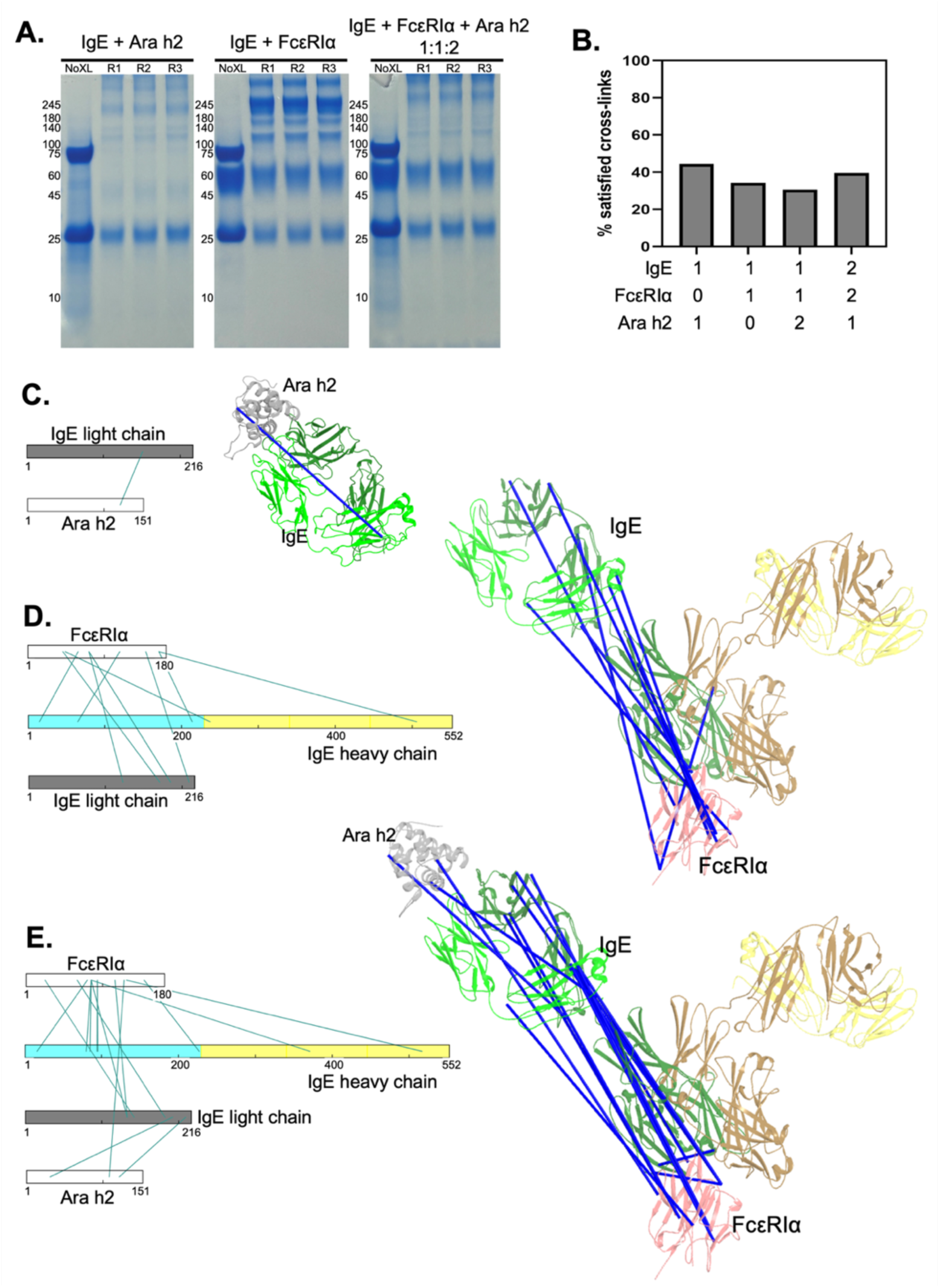
Interactions between IgE, FcεRIα, and Ara h 2 observed in XLMS. (**A**) Cross-linked protein complexes were analyzed using a stained SDS-PAGE gel. High molecular weight bands in cross-linked samples, compared to the no cross-link (NoXL) control, indicate the formation of covalent complexes. Molecular weight markers are indicated in kDa. (**B**) Percentage of cross-links that satisfy expected antibody-antigen distance constraints in cross-linked samples of IgE-Ara h 2, IgE-FcεRIa, IgE-FcεRIα-[Ara h 2]_2_ (1:1:2), and [IgE]_2_-FcεRIα]_2_-Ara h 2 (2:2:1). (**C-E**) Inter-protein cross-links for IgE-Ara h 2, IgE-FcεRIα, IgE-FcεRIα-Ara h 2 (1:1:2), respectively, are mapped onto the protein sequences (left) and structures (right), in which blue lines represent the cross-links.

Most cross-links were compatible with expected Cα-Cα distance constraints (Figure 3B), supporting consistency with the structural models. In total, 37 unique inter-links and 96 intra-links were identified. IgE-Ara h 2 complex showed fewer inter-links than the FcεRIα-containing assemblies, whereas intra-links within Ara h 2 were observed, particularly in helical regions, indicating an internally constrained allergen structure (Table S2). Overall, these data reveal non-canonical IgE binding modes with unexpected contacts between IgE and its binding partners.

In IgE-Ara h 2 complexes, an inter-link between Ara h 2 K133 (near region-III) and IgE light chain K151 (Cκ), shown in Figure 3C, affirmed IgE interaction near region-III epitope, aligning with both simulations and HDX-MS derived findings. This K133 interaction resulted in stabilization of residues 112-132 (*blue* in Figure 2C) which may be coupled with increased flexibility in the adjacent residues 133-145 (*red* in Figure 2C). This contact likely underlies the transient region-III engagement observed in simulations (Figure S7).

Additional IgE-Ara h 2 inter-links were detected in the presence of FcεRIα, including Ara h 2 K43 (region-II) paired with IgE light chain K192 (Cκ), and Ara h 2 S26 (region-I) paired with IgE heavy chain K43 (framework supporting hCDR1 and hCDR2), depicted in Figure 3E and S7, respectively. The detection of cross-links across all three epitope regions supports that IgE engages multiple Ara h 2 epitopes, albeit away from canonical paratopes. These observations imply that Ara h 2 can partially disengage from the IgE CDRs (consistent with BLI measurements in Figure S9), while remaining proximal to Fab. Notably, the greater number and diversity of cross-links IgE-Ara h 2 in the presence of FcεRIα suggest that FcεRIα binding may stabilize alternative or extended IgE-Ara h 2 interaction modes.

Unexpected cross-links between FcεRIα Y80 and IgE Fabs, alongside expected contacts with the FcεRIα-binding region (Cε2–Cε3), in Figure 3D and Table S2, suggests a subpopulation of IgE conformations, where Fabs approach FcεRIα, consistent with certain previously proposed IgE models^18,41^. Additionally, cross-links between FcεRIα S85 and IgE S371 (Cε3), detected only in the presence of Ara h 2, implies antigen-dependent stabilization of specific IgE-FcεRIα interfaces. Together, these results support a model in which Ara h 2 engagement promotes a more “closed” IgE Fc conformation, resembling the FcεRIα-bound state.

### Allosteric communication within IgE promoted by binding of Arah2 and/or FcεRIα

To investigate causal relationships governing IgE conformational changes, directed acyclic graphs (DAGs) were constructed for each IgE state, in which nodes represent structural descriptors and edges represent inferred dependencies. These graphs enabled causal inference of domain movements and potential allosteric pathways within IgE.

Edge weights (ω ∈ [0,1]) were derived from “parent importance” values obtained from a fitted non-linear SEM (Figure 4A). Across all states, weighted DAGs revealed strong directional dependencies, especially involving the FcεRIα-binding region (highlighted in red box). The edge weights were distributed heterogeneously, with a subset of highly weighted edges (ω ≥ 0.8) indicating dominant interdomain relationships that primarily drive conformational variance, particularly more pronounced in the bent states. These findings suggest IgE dynamics appear hierarchically propagated through selective structural constraints.

**Figure 4:**
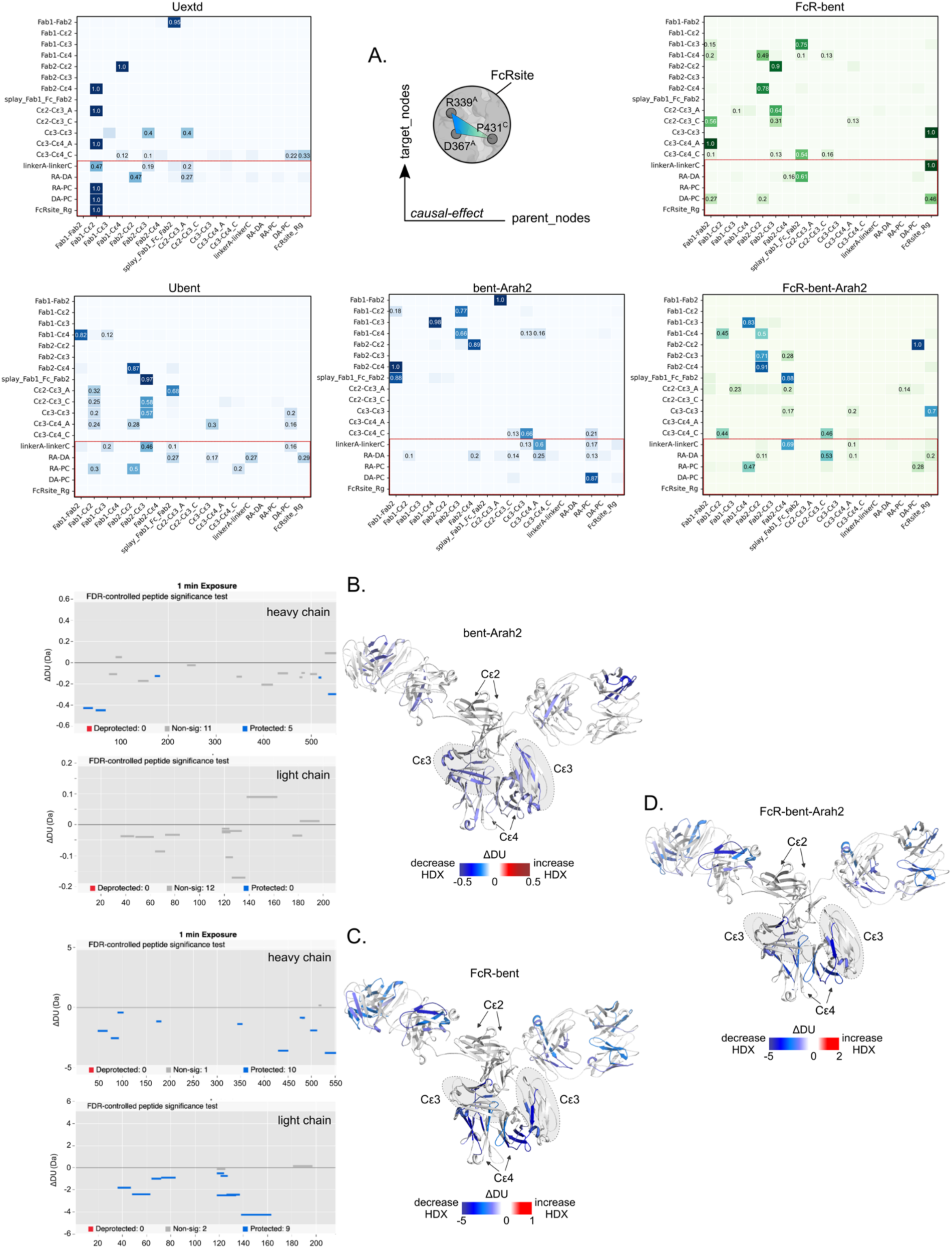
Possible allosteric communications within IgE structures when engaging with Ara h 2 and/or FcεRIα. **(A)** Causal graphs demonstrating effects of domain movements are presented as heatmaps with “parent importance” values used as weights (*w* ∈ [0,1]) for each IgE state. For clarity, only those edges carrying values ≥ 0.1 were annotated. **(B-D)** Woods plot showing differences in deuterium exchange (ΔDU) calculated for the Ara h 2-bound IgE state (*bent-Arah2*) relative to the free IgE state (B), and for the FcεRIα-bound IgE (*FcR-bent*) state relative to the free IgE state (C) at 1-min deuterium labelling time. The length of each line represents the corresponding pepsin proteolyzed peptide, shown along x-axis from N- to C-termini. A 99% confidence interval (CI) was used to analyze each labelling time. Peptides showing differences greater than CI are highlighted as per color key. Corresponding differences in deuterium uptake were mapped on to the full-length IgE structure for 1 min labelling time under different binding conditions: *bentArah2* (B), *FcR-bent* (C), and concurrent Ara h 2 and FcεRIα binding, *FcR-bent-Arah2* (D). For clarity, Ara h 2 and FcεRIα structures were excluded.

Comparing *Uextd* and *Ubent* states revealed distinct causal architectures. The *Ubent* network exhibited broader connectivity, demonstrating broader propagation of conformational variance that reflect greater allosteric connectivity. In contrast, the *Uextd* network was sparser with stronger dependencies, reflecting greater structural constraints. Relative to *Ubent*, both *bent-Arah2* and *FcR-bent* showed reduced network density, suggesting that Ara h 2 or FcεRIα binding independently restricts and refocuses causal constraints through limited dominant pathways. Particularly in *FcR-bent*, the variance was concentrated within specific interactions, involving Fabs-Fc splay angles, Fab1-Fab2, Fab2-Cε2, and Fab2-Cε4 that influence the FcεRIα-binding region. Overall, IgE dynamics are governed by a few critical interdomain interactions that disproportionately regulates the FcεRIα-binding region complexity.

HDX-MS supports these computational findings. In the *bent-Arah2*, reduced deuterium uptake across multiple regions, especially along the heavy chain, indicated global stabilization. As expected, heavy chain CDRs (residues 21-41 and 47-68) were protected, confirming the Ara h 2 engagement (Figure 4B and S12). While peptides spanning Cε4 (residues 515-521 and 528-552) showed early protection (1min), peptides spanning Cε3 (residues 395-419) exhibited decreased uptake at longer labeling times (Figure S10B-C, and S12), suggesting that Ara h 2 binding stabilizes IgE, potentially priming receptor-interacting regions.

In the *FcR-bent*, deuterium exchange was decreased across both heavy and light chains, including protection at FcεRIα-binding hotspot (residues 343-354) and the Cε3 DE loop (residues 429-451), confirming the FcεRIα engagement^17^. In addition, protection across multiple peptides spanning Cε4 (residues 429-451, 475-481, 479-486, 497-512, 515-521, and 528-552) indicated stabilization of Cε4-Cε4 interactions (Figure 4C and S12). Interestingly, reduced exchange within Fab regions suggested that FcεRIα binding might induce a more compact structure with decreased solvent exposure (Figure 4C and S13). Similarly, the tertiary Ara h 2-IgE-FcεRIα complex (*FcR-bent-Arah2*) showed decreased exchange across Cε3 and Cε4 along with CDRs region, confirming concurrent Ara h 2 and FcεRIα bindings (Figure 4D and S14). Overall, these results indicate that FcεRIα binding propagated long-ranged conformational stabilization, promoting a more compact bent conformation.

The *FcR-bent-Arah2* network exhibited sparse and “hub-dominated” topology with fewer yet stronger interdomain dependencies, involving mostly Fab2 and Cε2/Cε3/Cε4 domains (Figure 4A). This suggests that FcεRIα engagement may precondition IgE for subsequent Ara h 2 binding. The FcεRIα and Ara h 2 interplay reflects a distinct and structurally constrained state of IgE, where the presence of both partners may increase the Fab-Fc communications. For instance, Ara h 2 binding at Fab1 may influence Fab2 mobility towards Fc domains (Cε2, Cε3, and Cε4), consequently influencing the FcεRIα-binding site. Such Fab2-Fc proximal was also supported by XLMS (Figure 3D-E). Further, the allosteric communication within IgE appears to be reconfigured, shifting the dominant pathway from Fab2-Cε3 (*Ubent*) to Fab2-Cε4 upon the sequential binding, implying that different Cε domains mediate structural signaling in a state-dependent manner.

In brief, FcεRIα engagement may prime IgE for Ara h 2 recognition, while Ara h 2 binding further stabilizes Fabs and, in turn, bias IgE toward FcεRIα-bound conformations competent for downstream signaling. These responses, however, remain limited to the sampled conformations, DAG assumptions, and HDX-MS limitations, and thus the inferred mechanistic insights warrant further validation.

## DISCUSSION

We characterized full-length IgE dynamics in complexes with Ara h 2 and/or FcεRIα to address limitations of existing data derived from isolated domain structures. By combining MD simulations, XLMS, causal graph analysis, and HDX-MS, we examined domain proximities and inferred allosteric communications within the Ara h 2-IgE-FcεRIα system.

Causal network analysis revealed state-dependent IgE architectures. The unbound bent state showed broader yet weaker connectivity, whereas the unbound extended states exhibited more constrained dependencies. While this pattern is physically plausible, whether this difference reflects genuine distinct conformational reorganization or instead arises from complexity limitations of the IgE system, remains inconclusive. Nevertheless, these analyses suggest that a few dominant interdomain interactions govern IgE structural dynamics. A notable finding in the Ara h 2-bound state is the strong causal link between Fab-Fc coupling, particularly involving Fab2 and Cε domains, suggesting synergistic motions of Fabs that constrain the FcεRIα-binding region. HDX-MS data partially corroborates this, showing increased Fc stabilization. However, the concurrent protection of Cε3 indicates broader Fc constraints beyond Cε4 alone, which is not fully captured by the present causal analysis.

Upon FcεRIα engagement, dominant causal communication shifted from Cε3 to Cε4, aligning with HDX-MS data, with persistent Cε4 protection but diminished Cε3 protection relative to the Ara h 2-bound state alone (Figure 4C-D vs 4B). This supports a hypothesis that FcεRIα binding selectively stabilizes Cε4 while redistributing the restriction away from Cε3. While the HDX-MS observations are supportive, they do not constitute direct evidence for the specific allosteric pathways proposed by the DAGs. Thus, despite encouraging, these insights remain complementary rather than definitive mechanistic evidence.

Unlike IgG, IgE lacks a flexible hinge region, and instead contains an additional Cε2 domain^42^, resulting in restricted Fab flexibility and unique conformational dynamics^18,43^. Despite this, our findings show that the observed Fab flexibility may arise from geometric constraints imposed by the Cε2-Cε3-Cε4 architecture, representing a more limited yet functionally relevant motion. In fact, the Fab-Fc communication shifts from Cε3 to Cε4 upon concurrent Ara h 2 and FcεRIα binding, suggesting state-dependent Fab-mediated routing of conformational signals through the Fc. Taken together, these observations highlight unique conformational dynamics of IgE compared to IgG, where geometric constraints within the Fc region modulate, rather than abolish, Fab plasticity. These findings could have broader implications for understanding how Ara h 2-mediated crosslinking of FcεRIα-bound IgE is structurally propagated through the receptor-binding interfaces to initiate the allergic signaling.

Although this study characterizes a preliminary and hypothesis-generating conformational signature of Ara h 2 and/or FcεRIα engagement rather than a statistically confirmed structural transition in IgE, it provides a more holistic structural framework for Ara h 2 and/or FcεRIα-bound IgE complexes, offering new insights into IgE allostery and interdomain communication. As a complete structural model for the Ara h 2-IgE-FcεRIα complex is not yet available, this interdisciplinary approach provides a means to overcome the obstacles that impede the determination of its structure. Together, these findings may advance our understanding of peanut allergy and facilitate future mechanistic studies of allergic diseases in general.

## Abbreviations

IgE: Immunoglobulin-E
Ara h 2: *Arachis hypogaea* allergen 2
FcεRIα: high-affinity IgE receptor, Fc epsilon RI α-subunit
MD: Molecular Dynamics
XLMS: Cross-linking Mass Spectrometry
HDX-MS: Hydrogen-Deuterium Exchange Mass Spectrometry
SEM: Structural Equation Model
DAG: Directed Acyclic Graph
Uextd: unbound extended IgE
Ubent: unbound bent IgE
bent-Arah2: Ara h 2-bound bent IgE
FcR-bent: FcεRIα-bound bent IgE
FcR-bent-Arah2: concurrent Ara h 2 and FcεRIα-bound bent IgE

## Acknowledgments

We thank Waiheng Lua and Nir Kalisman for helpful discussions on IgE complexity in process of producing and purifying IgE as well as XLMS experiments, and thanks the Protein and Proteomics Centre (PPC) in the National University of Singapore for equipment supports in the HDX-MS experiments. MD simulations were performed on the ASPIRE-2A clusters of the National Supercomputing Center of Singapore. This work was supported by the National Medical Research Council grant NMRC-OFYIRG (MOH-000661) awarded to CTTS and partially by the A*STAR Young Achiever Award awarded to ZS. We also acknowledge BII (A*STAR) core funds.

The authors used Copilot, a built-in in the Microsoft 365 (Word) to assist in language improvement.

## Data availability

Raw cross-linking mass spectrometry spectra and search data have been deposited at ProteomeXchange^44^ (PXD076139) and JPost repository^45^ (JPST004513).

(For reviewers: https://repository.jpostdb.org/preview/148879281869c3a7917c49c, Access key 9634) HDX-MS details are included in the Supplementary File.

During the preparation of this work the author(s) used Copilot, a built-in in the Microsoft 365 (Word) in order to assist in language improvement. After using this tool/service, the author(s) reviewed and edited the content as needed and take(s) full responsibility for the content of the publication

