## Supplementary_File for "Ara h 2 and FcεRIα binding elicit state-dependent allostery in IgE"

**Supplementary Figures and Tables**

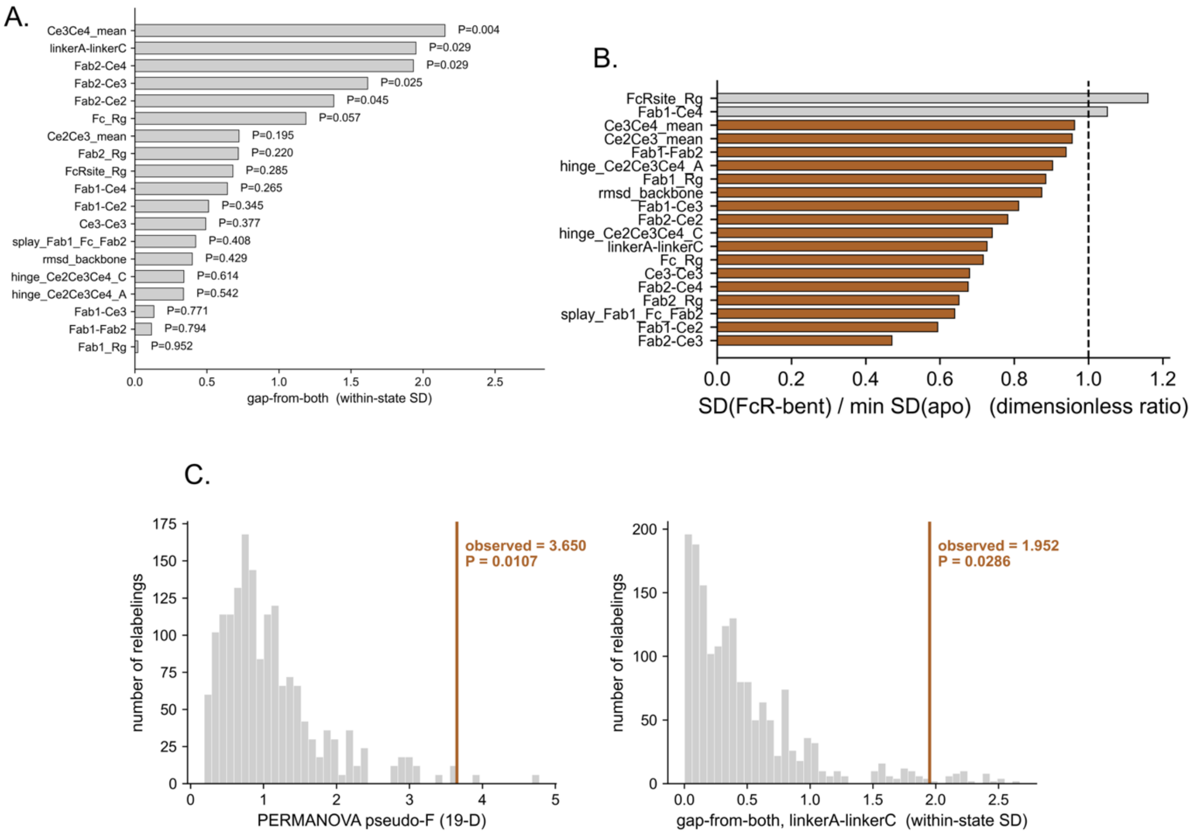


**Figure S1:** Statistical tests of 19 measurable descriptors of domain distances and angles for comparison across *Uextd, Ubent,* and *FcR-bent* states of IgE. **(A)** The 19 variables were ranked (highest on top) based on the separation of the *FcR-bent* ensemble from both the unbound states *Uextd* and *Ubent*. None of the variables individually remains significant after the Benjamini-Hochberg false discovery rate (FDR) correction, e.g. the best rank Cε3-Cε4_mean having uncorrected p-value *P*=0.004 (see details below) and corrected p-value *q*=0.068 **(B)** The *FcR-bent* shows less wobble than both the unbound states referenced on 17/19 measurements. **(C)** The statistical basis underlying every reported P-value. For example, given independent trajectories (n=3) per state, the omnibus three-state permutation test (1680 exact relabeling) and the linkerA-linkerC gap statistic are each referenced against their full null distributions (observed pseudo-F = 3.65, P = 0.0107; observed gap = 1.952 SD, P = 0.0286), with a baseline of 1/1680 (≈ 0.0006).

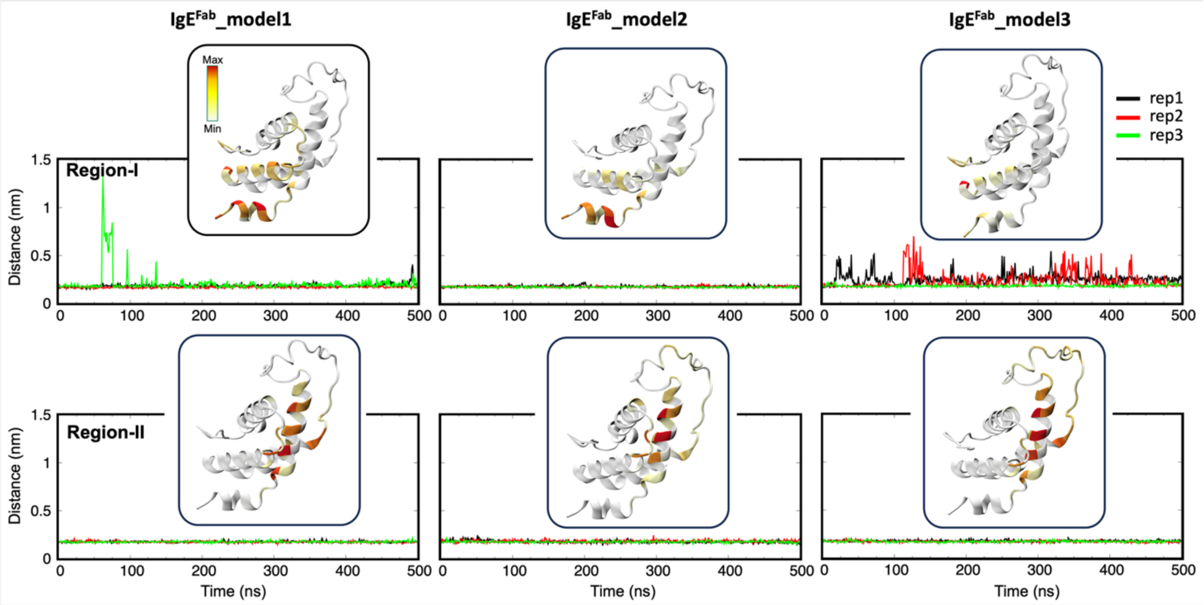


**Figure S2:** Interaction analysis of IgE-Ara h 2 complexes. The distances between the centers of mass of region-II and IgE-CDRs showed that binding is more stable to this region compared to those in region-I. Contact analysis was performed using "Interquant" (implemented in VMD, doi:10.1016/0263-7855(96)00018-5) and visualized using heatmap: the Ara h 2 residues are colored from white to red, indicating the percentage of simulation time during which the interactions occurred. The Interactions were defined within inter-atomic distances of < 4.0 Å.

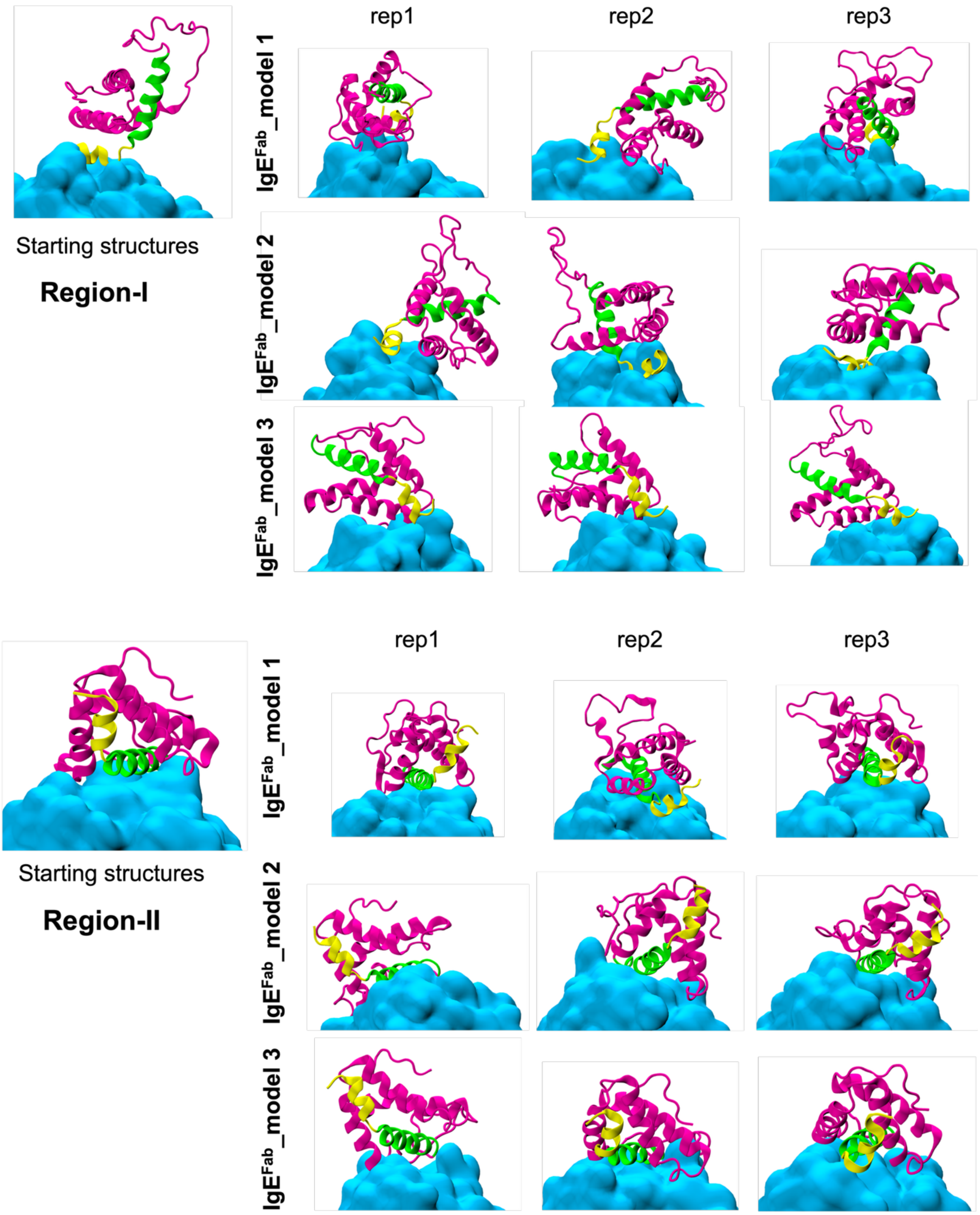


**Figure S3:** The two interfacial regions did not exhibit simultaneous contacts to the IgE-Fab1. Last frames of the simulations are presented with regions I and II in yellow and green cartoons, respectively while IgE-Fab1 is shown in cyan surfaces. For simplicity, only the interaction regions are highlighted.

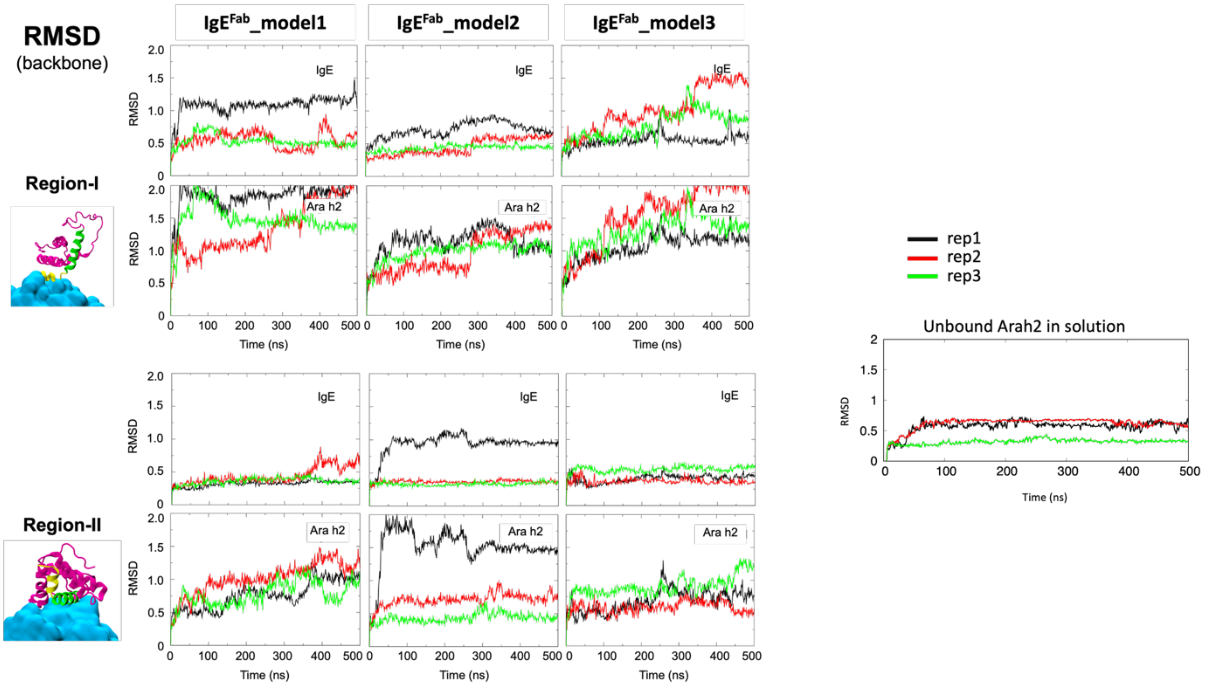


**Figure S4:** Calculated backbone RMSD of the IgE-Fab1 and Ara h 2 at the interface regions (left) and of the unbound Ara h 2 in solution (right).

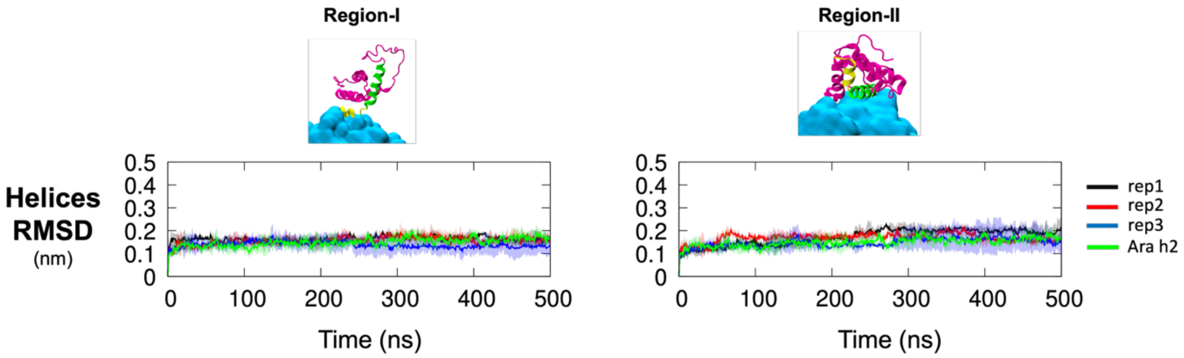


**Figure S5:** Calculated RMSD of helical regions within IgE-Fab1 and Ara h 2 at the two interface regions.

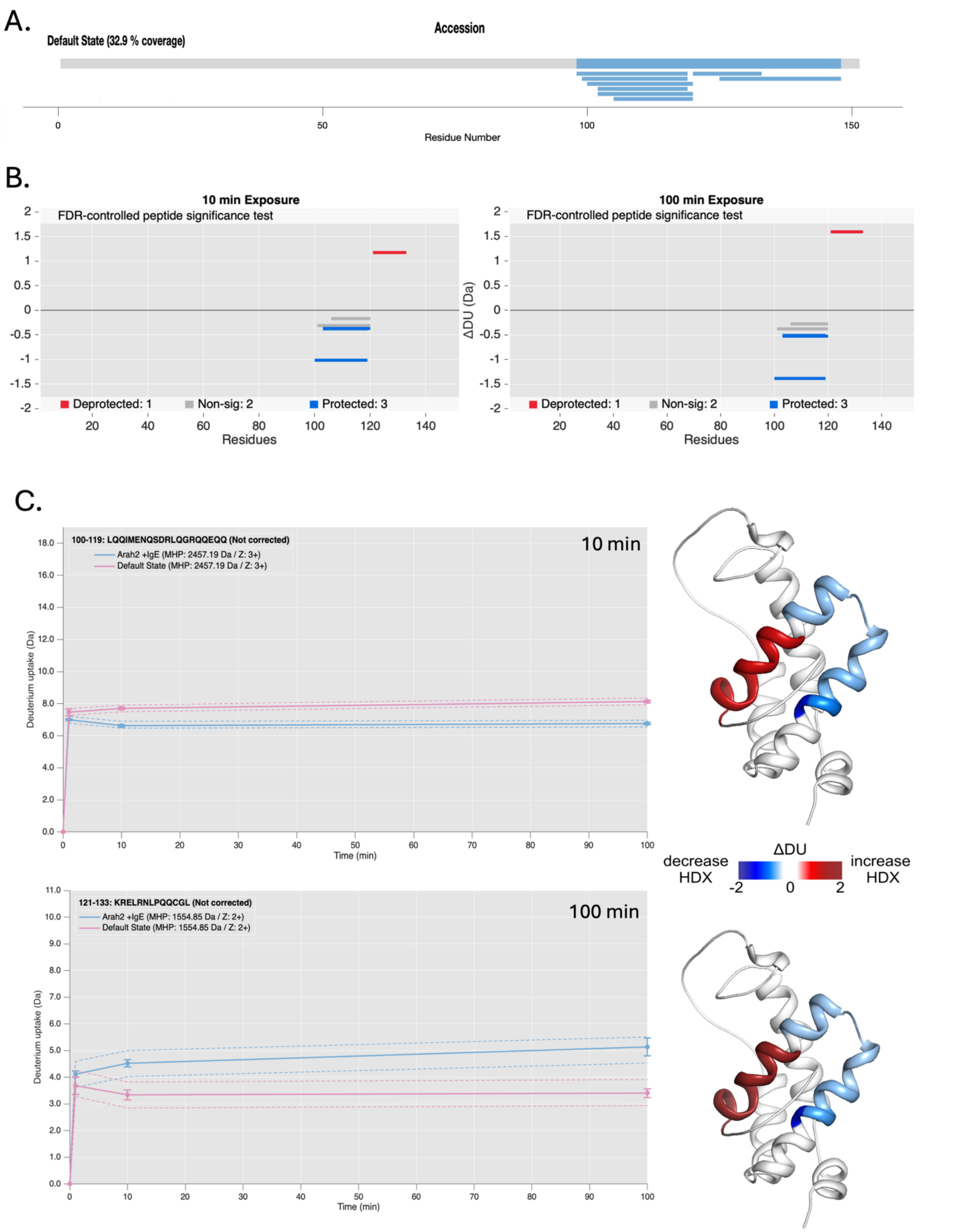


**Figure S6**: (A) Coverage map from pepsin proteolyzed peptide of Ara h 2 protein from HDX-MS experiments, with 32.9% sequence coverage. (B) Woods plot showing differences in deuterium exchange (ΔDU, Y-axis) between IgE-Ara h 2 complex state and free Ara h 2 state at deuterium labelling times, t = 10 and 100 min. The length of the lines represents the length of each pepsin proteolyzed peptides listed along X-axis from N- to C-termini. The confidence interval (CI) used to analyze each labelling time is 99%. Peptides showing differences greater than CI are highlighted as per color key. (C) Deuterium uptake plots showing peptide 100-119 and 121-133 (renumbered to 112-131 and 133-145 in the main text) with respect to time. (D) Differences in deuterium uptake between IgE-Ara h 2 complex and free Ara h 2 state mapped on to the Ara h 2 structure for labelling times of 10 and 100 minutes.

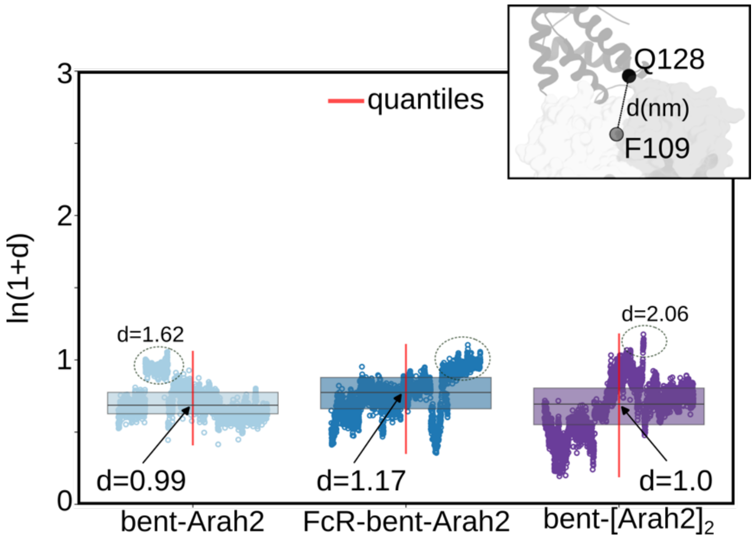


**Figure S7**: Distance between Ara h 2 and IgE-Fab1 binding region in various Ara h 2-bound simulations to demonstrate possible transient binding of Ara h 2 to IgE. Distances were measured using the two highlighted residues Q128 and F109 considered as center-of-mass of Ara h 2 (region-**III** epitope) and IgE-Fab1, respectively.

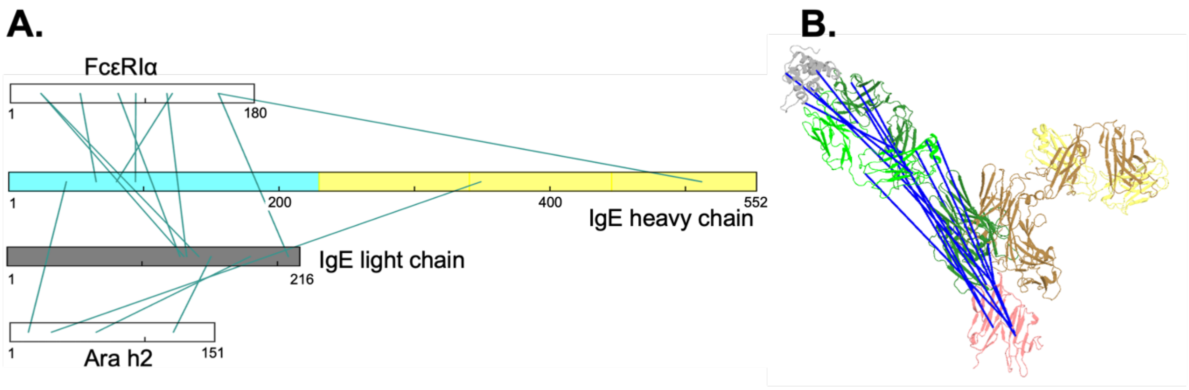


**Figure S8**: Cross-links of IgE-FcεRIα-Ara h 2 complexes (ratio 2:2:1). Inter-protein cross-links are represented on the protein sequences (A) and structures (B), in which lines represent the cross-links.

| Method | Ligand | Ligand Conc. (μg/ml) | Immobilization Level (nm) | Analyte | Analyte Conc. (nM) | Association time (s) | Dissociation time (s) |
|---|---|---|---|---|---|---|---|
| NTA | Arah1 | 2.5 | ~2.0 | IgE | 1000, 500, 250, 125 | 300 | 300 |
| | Arah2 | 0.4 | ~0.2 | | 100, 50, 25, 12.5, 6.25 | 120 | 300 |
| | FcERIa | 0.4 | ~0.2 | | 100, 50, 25, 12.5, 6.25 | 120 | 160 |

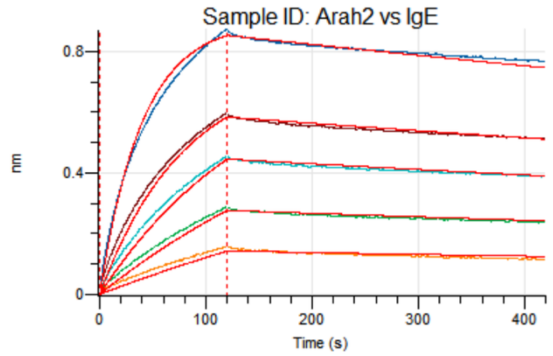


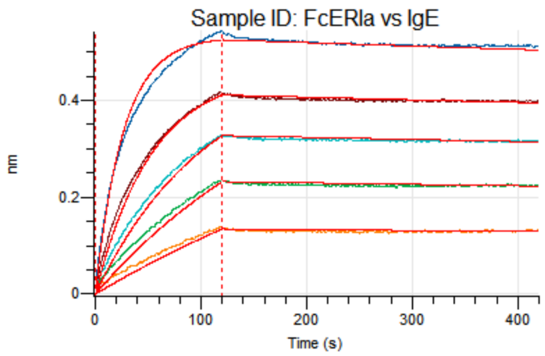


| Ligand | Analyte | Rmax | Ka (1/Ms) | Kdis (1/s) | KD (M) | Dissoc X^2 | Dissoc R^2 |
|---|---|---|---|---|---|---|---|
| FcERIa | IgE | 0.533 | 3.729E05 | 1.368E-04 | 3.667E-10 | 0.7067 | 0.9968 |
| Arah2 | | 0.9144 | 2.445E05 | 4.458E-04 | 1.823E-09 | 1.8452 | 0.9969 |

**Figure S9A**: Binding kinetics of Ara h 2 and FcεRIα to IgE, performed by GenScript USA Inc., using the Octet® BLI Discovery version 12.2.2.26. The assay was performed at 30°C and at 1000 rpm. Ara h 2 and FcεRIα protein were firstly immobilized onto NTA biosensor. IgE was applied as analyte for association and dissociation steps. Association and dissociation of Ara h 2 and FcεRIα interacting with IgE monitored by Octet$^{RED}$ 384. Ara h1 was also attempted in the binding tests but was found not to bind IgE (data not shown).

All the data were processed using the Octet$^{R}$ BLI Discovery version 12.2.2.26.

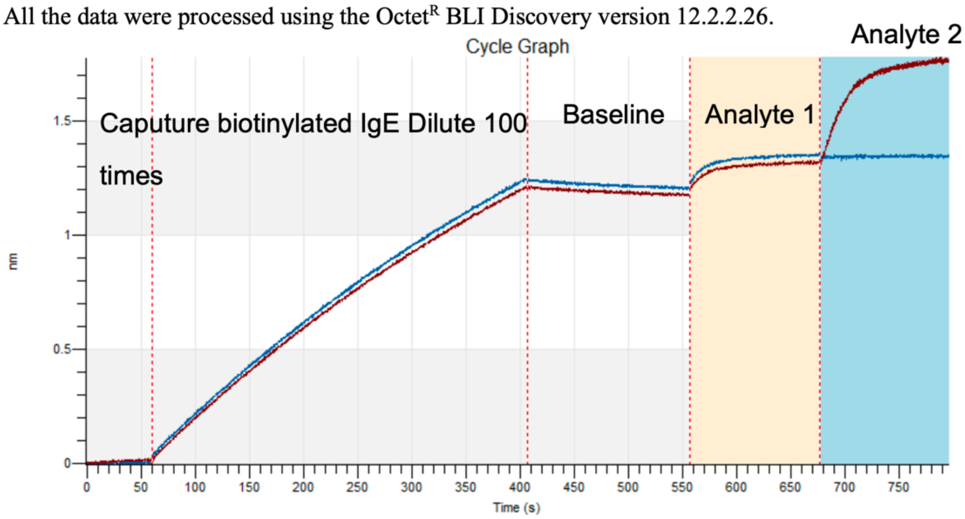


**Figure 1 Cycle Graph**

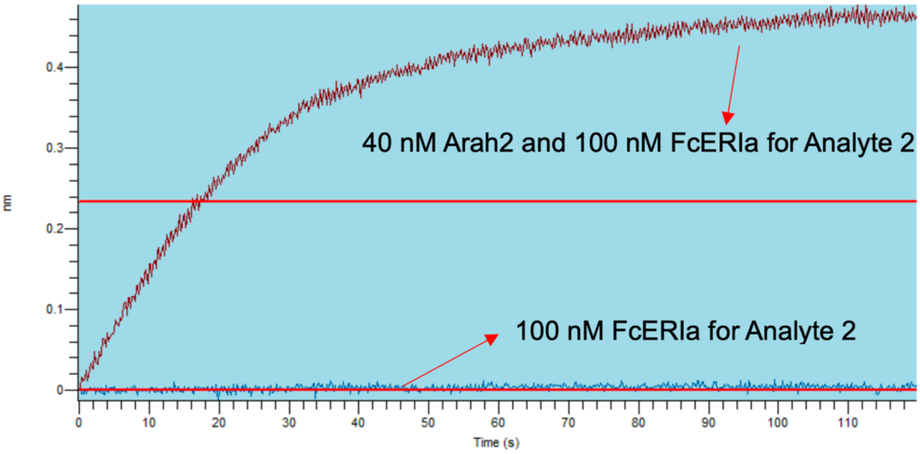


**Figure 2 Analyte 2 Step Graph**

| Info | Sample ID | Average signal (nm) |
|---|---|---|
| Analyte 1 | FcERIa | 0.124 |
| Analyte 2 | Arah2 | 0.4637 |
| Negative control | FcERIa | 0.0035 |

**Figure S9B**: Binding kinetics of triple bindings of Ara h 2 and FcεRIα to IgE, performed by GenScript USA Inc., using the Octet® BLI Discovery version 12.2.2.26. The assay was performed at 30°C and at 1000 rpm. Dilute 100 times biotinylated IgE was firstly immobilized onto SA biosensor with immobilization Level of ~0.13 nm. 100 nM FcεRIα and 40 nM Ara h 2 was applied as Analyte 1 and Analyte 2 for association steps. 100 nM FcεRIα was applied as Analyte 2 for negative control.

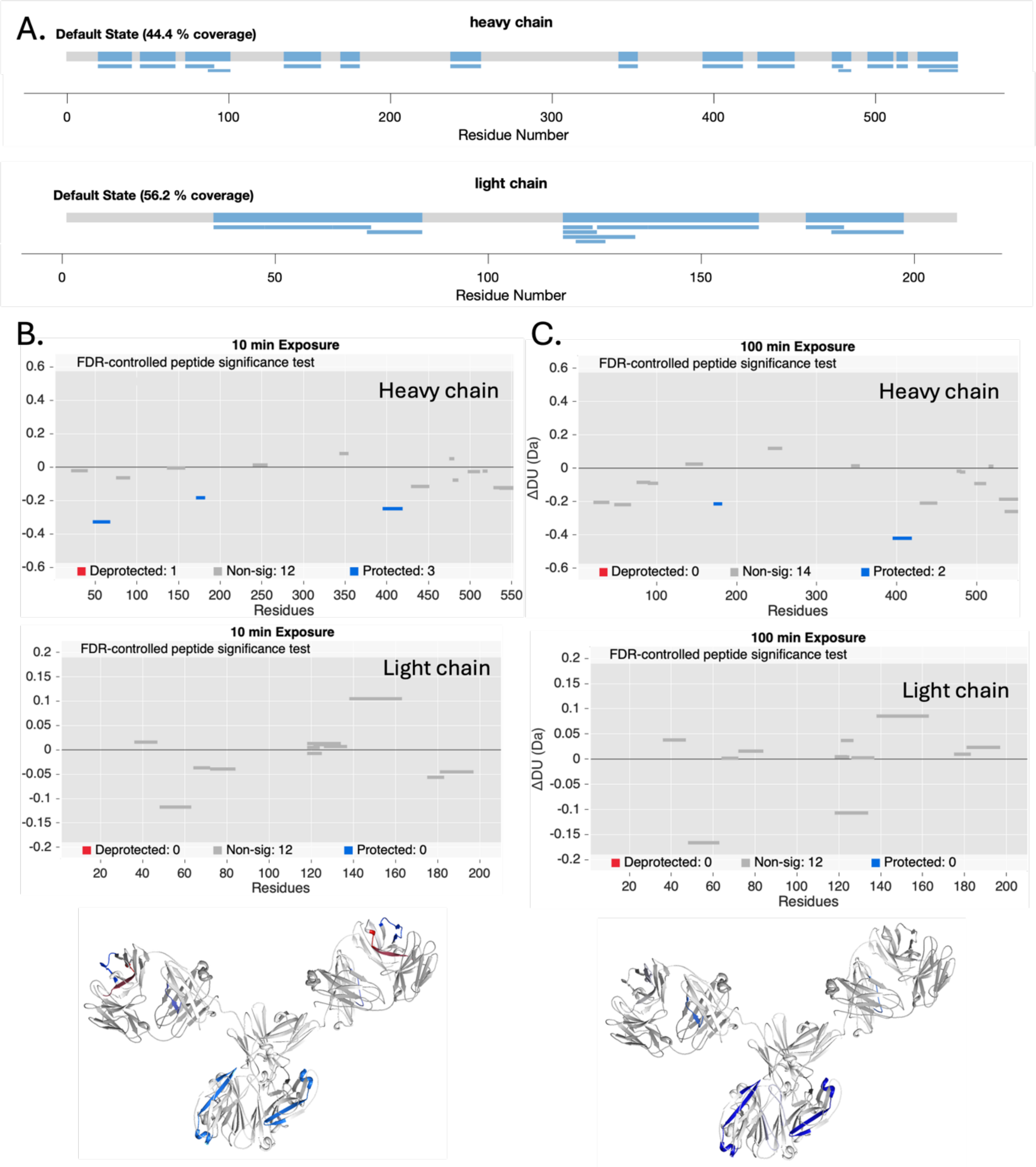
A.
heavy chain
Default State (44.4 % coverage)
0
100
200
300
400
500
Residue Number
light chain
Default State (56.2 % coverage)
0
50
100
150
200
Residue Number
B.
10 min Exposure
FDR-controlled peptide significance test
Heavy chain
Deprotected: 1
Non-sig: 12
Protected: 3
Residues
10 min Exposure
FDR-controlled peptide significance test
Light chain
Deprotected: 0
Non-sig: 12
Protected: 0
Residues
C.
100 min Exposure
FDR-controlled peptide significance test
Heavy chain
ΔDU (Da)
Deprotected: 0
Non-sig: 14
Protected: 2
Residues
100 min Exposure
FDR-controlled peptide significance test
Light chain
ΔDU (Da)
Deprotected: 0
Non-sig: 12
Protected: 0
Residues

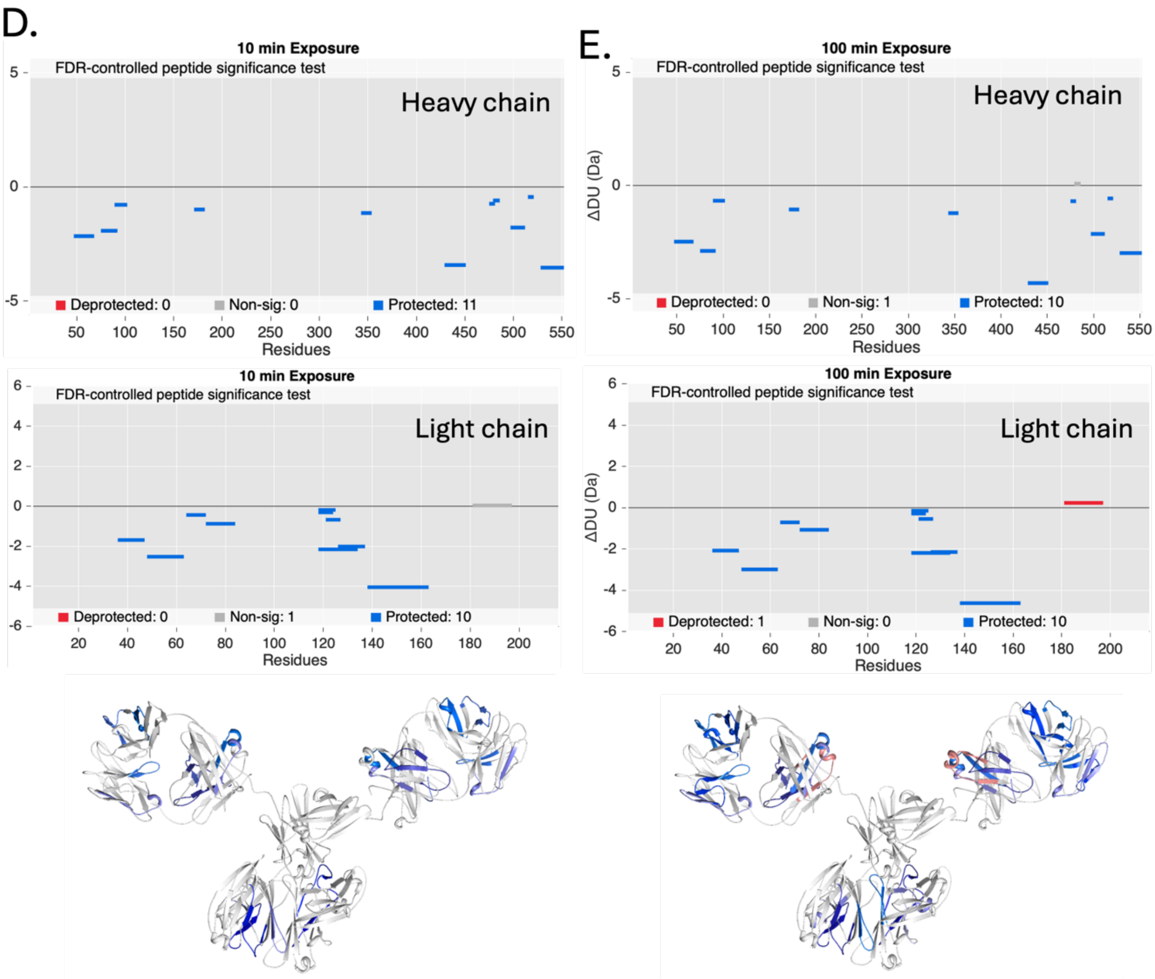


**Figure S10**: **(A)** Coverage map from pepsin proteolyzed heavy and light chain IgE peptides from HDX-MS experiments, with 44.4% and 56.2% sequence coverage, respectively. **(B-E)** Woods plot showing differences in deuterium exchange (ΔDU, Y-axis) within IgE estimated from the Ara h 2-bound IgE state against the free IgE state (B and C) and within IgE estimated from the FcεRIα-bound IgE state against the free IgE state (D and E) accordingly at 10 min and 100 min deuterium labelling time. The length of the lines represents the length of each pepsin proteolyzed peptides listed along X-axis from N to C-termini. Differences in deuterium uptake were mapped on to the full-length IgE structure accordingly for labelling times 10 min and 100 min. For better visualization, Ara h 2 and FcεRIα structures were excluded.

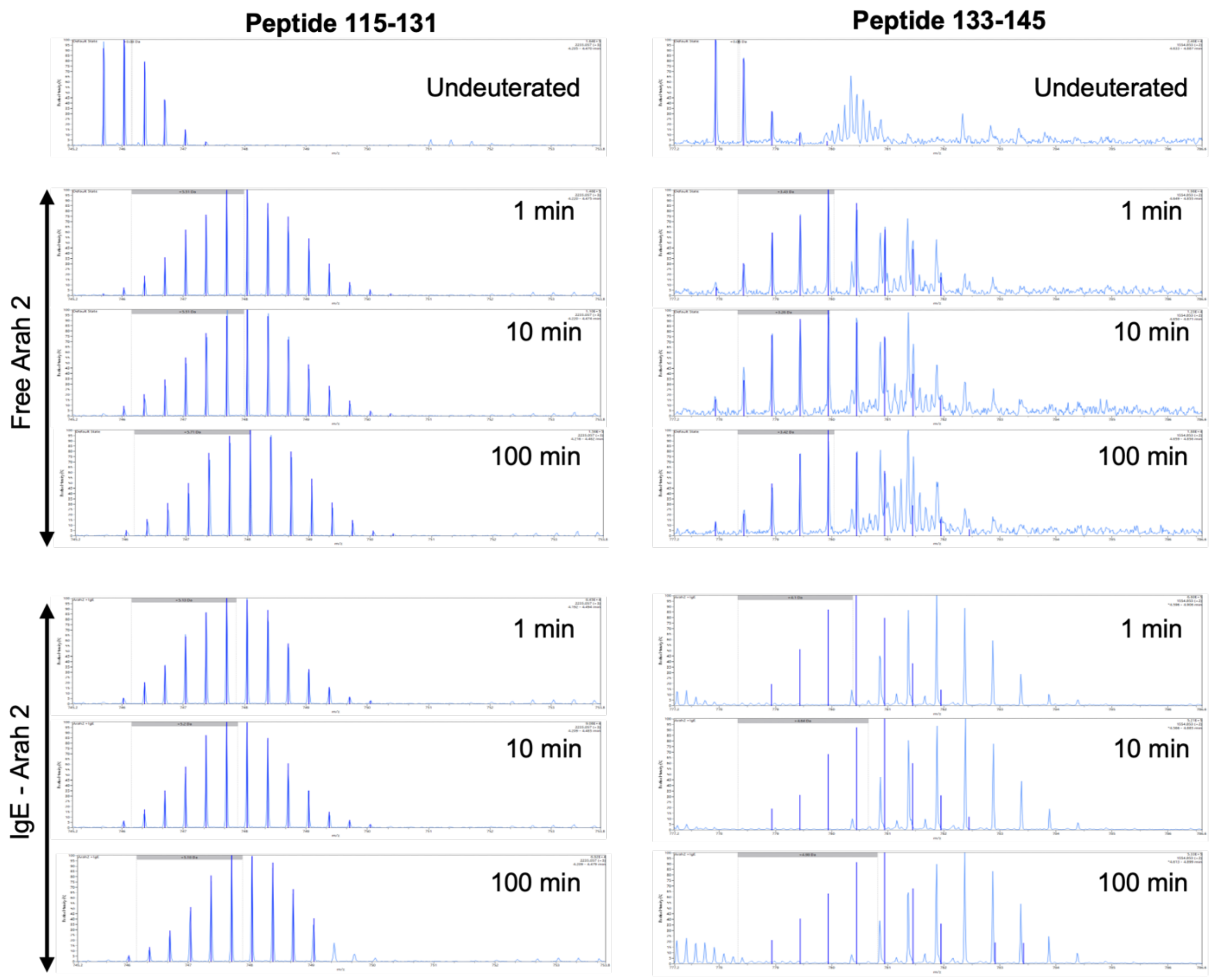


**Figure S11**: Mass spectra with isotopic envelopes after deuterium exchange (t = 1, 10 and 100 min) for selected peptide 115-131 (left) and peptide 133-145 (right) from Ara h 2 shown for the free Ara h 2 and Ara h 2-IgE complex. Mass spectra of the equivalent un-deuterated peptide is shown for reference. The centroid masses are indicated label and dotted line.

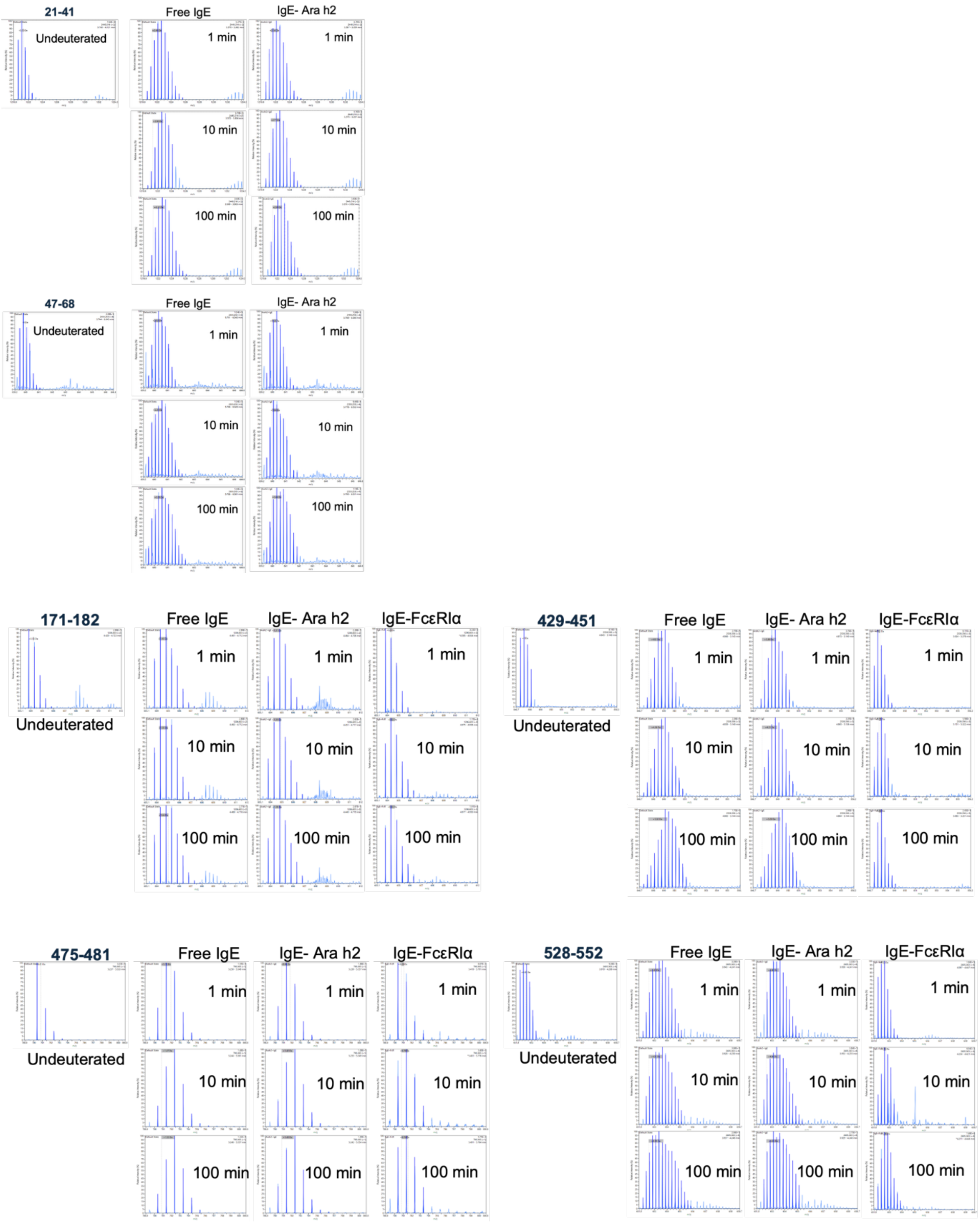


**Figure S12**: Mass spectra with isotopic envelopes after deuterium exchange (t = 1, 10 and 100 min) for select peptides of IgE heavy chain, shown for the free IgE, IgE-Ara h 2 and IgE-FcεRIα complexes. Mass spectra of the equivalent un-deuterated peptide is shown for reference. The centroid masses are indicated label and dotted line.

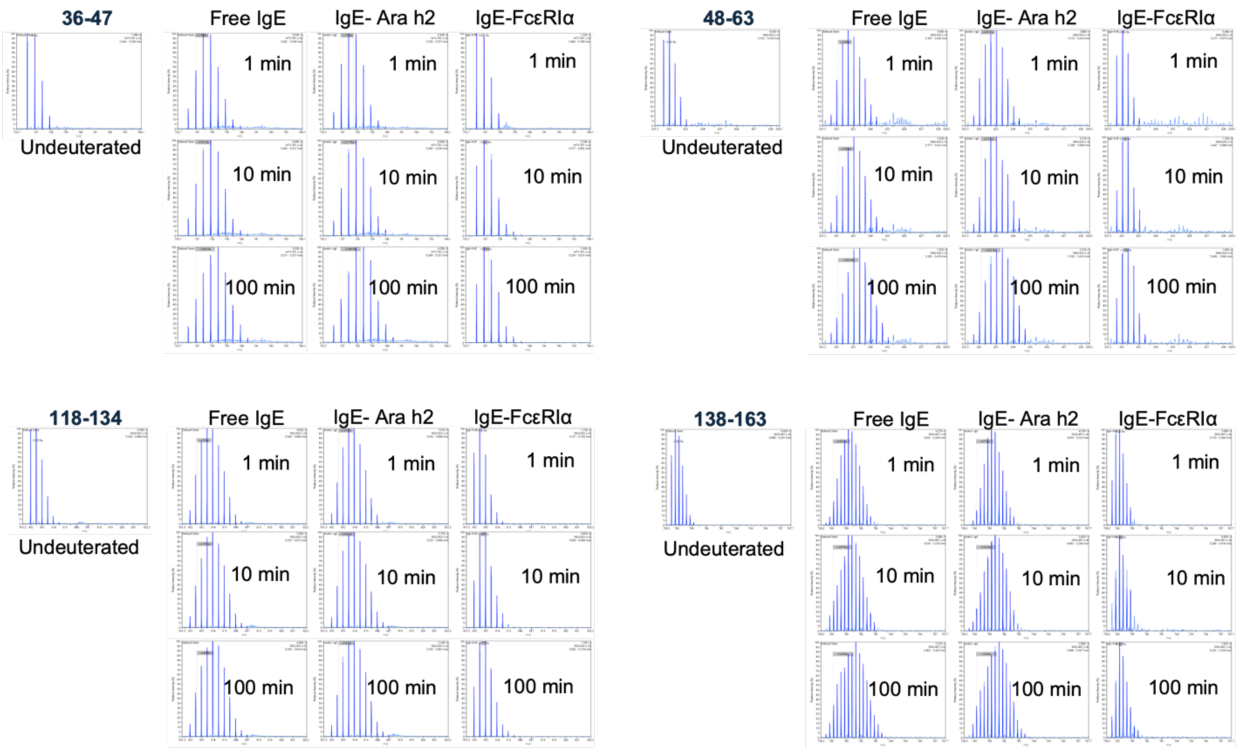


**Figure S13**: Mass spectra with isotopic envelopes after deuterium exchange (t = 1, 10 and 100 min) for select peptides of IgE light chain, shown for the free IgE, IgE-Ara h 2 and IgE-FcεRIα complexes. Mass spectra of the equivalent un-deuterated peptide is shown for reference. The centroid masses are indicated label and dotted line.

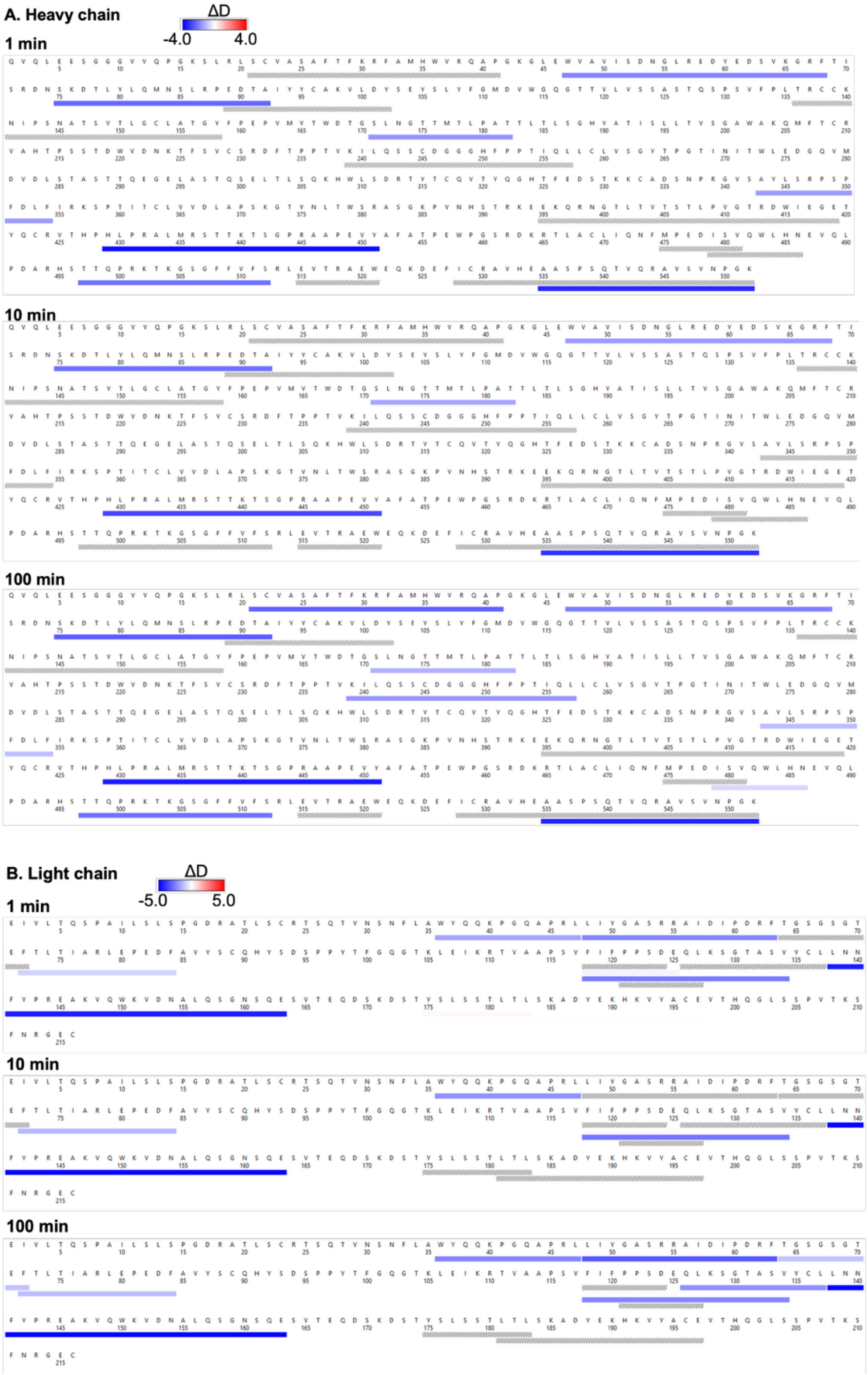


**Figure S14**: Deuterium difference between the tertiary Ara h 2-IgE-FcεRIα complex and the free IgE. Heat map showing differences in deuterium exchange at deuterium labelling times: t = 1, 10 and 100 min for heavy chain (A) and light chain (B) along with sequences and lines present the length of peptide and the difference in deuterium uptake for each pepsin protolyzed peptide are color-coded as per key.

**Table S1:** Binding affinity (kcal/mol) of $IgE^{Fab1}$ and Ara h 2.

| | **Ara h 2-$IgE^{Fab1}$_model1** | **Ara h 2-$IgE^{Fab1}$_model2** | **Ara h 2-$IgE^{Fab1}$_model3** |
|---|---|---|---|
| **Region-I** | -8.73 ± 0.5 | -11.25 ± 0.29 | -9.03 ± 0.29 |
| **Region-II** | -9.32 ± 0.44 | -8.62 ± 0.95 | -8.28 ± 0.4 |

**Table S2**: Identified inter-links of the IgE complexes (HC: Heavy chain, LC: Light chain). The Ara h 2 sequence was renumbered according to Mueller *et al.* (PDB: 3OB4). Note: Ara h 2 residues are renumbered to be consistent with the computational models.

| Protein1 | Site1 | Protein2 | Site2 | Score | aa1 | Peptide 1 | aa2 | Peptide2 | Count |
|---|---|---|---|---|---|---|---|---|---|
| **IgE-Ara h 2** | | | | | | | | | |
| Ara h 2 | 118 | IgE_LC | 151 | 3 | K | NQSDRLQGRQQEQQFK | K | VQWKVDNALQSGNSQESVTE | 1 |
| **IgE-FcεRIα** | | | | | | | | | |
| IgE_HC | 15 | FcεRIα | 66 | 3 | K | QVQLEESGGGVVQPGK | Y | SGEYKBQHQQVNE | 1 |
| FcεRIα | 46 | IgE_LC | 171 | 3 | S | VSSTKWFHNGSLSEE | K | SVTEQDSKD | 1 |
| FcεRIα | 49 | IgE_HC | 237 | 2 | T | TNSSLNIVNAKFEDSGE | T | WVDNKTFSVBSRDFTPPTVK | 1 |
| IgE_HC | 65 | FcεRIα | 120 | 4 | K | YEDSVK | Y | VIYYKDGEALK | 1 |
| FcεRIα | 80 | IgE_LC | 123 | 2 | Y | PVYLEVFSDWLLLQASAE | S | RTVAAPSVFIFPPSDEQLK | 2 |
| FcεRIα | 80 | IgE_LC | 185 | 3 | Y | PVYLEVFSDWLLLQASAE | K | STYSLSSTLTLSKADYE | 1 |
| FcεRIα | 154 | IgE_LC | 209 | 3 | K | SGTYYBTGKVWQLD | K | KHKVYABEVTHQGLSSPVTK | 1 |
| FcεRIα | 171 | IgE_HC | 214 | 2 | K | SEPLNITVIKAPRE | T | QmFTBRVAHTPSSTDWVDNK | 1 |
| FcεRIα | 171 | IgE_HC | 506 | 5 | K | PLNITVIK | S | TKGSGFFVFSRLEVTRAE | 1 |
| **IgE-FcεRIα-Ara h 2_112** | | | | | | | | | |
| IgE_HC | 16 | FcεRIα | 80 | 13 | K | ESGGGVVQPGK | Y | YKBQHQQVNESEPVYLE | 1 |
| FcεRIα | 25 | IgE_LC | 142 | 4 | T | GENVTLTBNGNNFFEVSSTK | Y | SGTASVVBLLNNFYPREAK | 1 |
| Ara h 2 | 38 | IgE_LC | 192 | 27 | K | QHLMQKIQRD | K | YEKHK | 1 |
| FcεRIα | 67 | IgE_LC | 182 | 3 | K | SGEYKBQHQQVNESE | T | STYSLSSTLTLSKADYEK | 1 |
| IgE_HC | 80 | FcεRIα | 85 | 7 | Y | NSKDTLYLQmNSLRPE | S | PVYLEVFSDWLLLQASAE | 1 |
| IgE_HC | 85 | FcεRIα | 85 | 7 | S | NSKDTLYLQmNSLRPE | S | PVYLEVFSDWLLLQASAE | 1 |
| FcεRIα | 85 | IgE_LC | 133 | 3 | S | VFSDWLLLQASAEVVmE | S | SGTASVVBLLNNFYPREAK | 1 |
| FcεRIα | 85 | IgE_HC | 371 | 6 | S | VFSDWLLLQASAEVVME | S | LAPSK | 1 |
| FcεRIα | 93 | IgE_HC | 94 | 18 | S | SEPVYLEVFSDWLLLQASAE | Y | TAIYYBAK | 1 |
| FcεRIα | 93 | IgE_HC | 95 | 8 | S | SEPVYLEVFSDWLLLQASAE | Y | TAIYYBAK | 1 |
| FcεRIα | 116 | IgE_LC | 133 | 12 | Y | VYKVIYYK | S | SGTASVVBLLNNFYPRE | 2 |
| Ara h 2 | 130 | IgE_LC | 209 | 41 | K | QQFKRE | K | VTHQGLSSPVTKSFNRGEB} | 1 |

| | | | | | | | | | |
|---|---|---|---|---|---|---|---|---|---|
| FcεRIα | 131 | IgE_HC | 517 | 3 | Y | YWYENHNISITNATVED | T | GSGFFVFSRLEVTRAEWE | 1 |
| FcεRIα | 154 | IgE_HC | 227 | 3 | K | SGTYYBTGKVWQLDYE | S | NKTFSVBSRDFTPPTVK | 1 |
| **IgE-FcεRIα-Ara h 2_221** | | | | | | | | | |
| Ara h 2 | 22 | IgE_HC | 43 | 3 | S | RRBQSQLERANLRPBE | K | RFAmHWVRQAPGKGLE | 1 |
| FcεRIα | 23 | IgE_LC | 131 | 9 | T | GENVTLTBNGNNFFEVSSTK | T | SGTASVVBLLNNFYPREAK | 1 |
| FcεRIα | 23 | IgE_LC | 142 | 8 | T | GENVTLTBNGNNFFEVSSTK | Y | SGTASVVBLLNNFYPREAK | 2 |
| Ara h 2 | 30 | IgE_HC | 349 | 4 | K | RANLRPBEQHLmQKIQRDE | S | SNPRGVSAYLSRPSPFD | 1 |
| FcεRIα | 52 | IgE_HC | 65 | 4 | S | ETNSSLNIVNAK | K | DSVKGRFTISRD | 1 |
| Ara h 2 | 64 | IgE_LC | 180 | 24 | S | RRDPYSPSPYD | T | STYSLSSTLTLSKADYE | 1 |
| IgE_HC | 80 | FcεRIα | 120 | 4 | Y | NSKDTLYLQMNSLRPE | Y | VYKVIYYK | 1 |
| FcεRIα | 80 | IgE_LC | 128 | 3 | Y | PVYLEVFSDWLLLQASAE | K | RTVAAPSVFIFPPSDEQLK | 1 |
| FcεRIα | 93 | IgE_HC | 94 | 23 | S | SEPVYLEVFSDWLLLQASAE | Y | TAIYYBAK | 1 |
| FcεRIα | 116 | IgE_LC | 133 | 5 | Y | VYKVIYYK | S | SGTASVVBLLNNFYPRE | 2 |
| Ara h 2 | 118 | IgE_LC | 151 | 8 | K | NQSDRLQGRQQEQQFK | K | VQWKVDNALQSGNSQESVTE | 1 |
| FcεRIα | 154 | IgE_LC | 208 | 4 | K | SGTYYBTGKVWQLD | T | KHKVYABEVTHQGLSSPVTK | 1 |
| FcεRIα | 154 | IgE_HC | 512 | 3 | K | DSGTYYBTGKVWQLD | S | GSGFFVFSRLEVTRAEWE | 1 |

**Table S3**: Identified IgE intra-links (HC: Heavy chain, LC: Light chain)

| Protein1 | Site1 | Protein2 | Site2 | Score | aa1 | Peptide 1 | aa2 | Peptide2 | Count |
|---|---|---|---|---|---|---|---|---|---|
| **IgE-Ara h 2** | | | | | | | | | |
| IgE_HC | 7 | IgE_HC | 21 | 16 | S | SGGGVVQPGK | S | SLRLSBVASAFTFK | 1 |
| IgE_HC | 52 | IgE_LC | 151 | 36 | S | GLEWVAVISDNGLRED | K | AKVQWKVDNALQSGNSQE | 1 |
| IgE_HC | 80 | IgE_LC | 111 | 5 | Y | NSKDTLYLQmNSLRPED | T | RTVAAPSVFIFPPSDE | 1 |
| IgE_HC | 80 | IgE_LC | 151 | 49 | Y | NSKDTLYLQMNSLRPE | K | AKVQWKVDNALQSGNSQE | 1 |
| IgE_HC | 85 | IgE_LC | 116 | 4 | S | DTLYLQmNSLRPE | S | IKRTVAAPSVFIFPPSD | 1 |
| IgE_HC | 85 | IgE_LC | 147 | 13 | S | NSKDTLYLQMNSLRPE | K | AKVQWKVDNALQSGNSQE | 2 |
| IgE_LC | 123 | IgE_LC | 128 | 81 | S | RTVAAPSVFIFPPSDE | K | QLKSGTASVVBLLNNFYPRE | 1 |
| IgE_LC | 123 | IgE_LC | 129 | 15 | S | RTVAAPSVFIFPPSDE | S | QLKSGTASVVBLLNNFYPRE | 1 |
| IgE_LC | 123 | IgE_LC | 133 | 27 | S | RTVAAPSVFIFPPSDE | S | QLKSGTASVVBLLNNFYPRE | 1 |
| IgE_LC | 123 | IgE_LC | 142 | 46 | S | RTVAAPSVFIFPPSDE | Y | QLKSGTASVVBLLNNFYPRE | 1 |
| IgE_LC | 151 | IgE_LC | 166 | 105 | K | VQWKVDNALQSGNSQE | T | SVTEQDSK | 1 |
| IgE_LC | 164 | IgE_LC | 185 | 45 | S | VDNALQSGNSQESVTE | K | QDSKDSTYSLSSTLTLSK | 2 |
| IgE_LC | 164 | IgE_HC | 234 | 15 | S | SVTEQDSK | T | NKTFSVBSRDFTPPTVK | 1 |
| IgE_LC | 166 | IgE_LC | 185 | 85 | T | VDNALQSGNSQESVTE | K | QDSKDSTYSLSSTLTLSK | 2 |
| IgE_LC | 199 | IgE_HC | 225 | 11 | T | VYABEVTHQGLSSPVTK | T | NKTFSVBSRDFTPPTVK | 1 |
| IgE_HC | 307 | IgE_HC | 314 | 32 | K | LTLSQKHWLSD | T | RTYTBQVTYQGHTFE | 1 |
| IgE_HC | 307 | IgE_HC | 325 | 6 | K | LASTQSELTLSQKHWLSD | T | RTYTBQVTYQGHTFEDSTK | 1 |
| IgE_HC | 552 | IgE_HC | 552 | 53 | K | AASPSQTVQRAVSVNPGK} | K | AASPSQTVQRAVSVNPGK} | 3 |
| **IgE-FcεRIα** | | | | | | | | | |
| IgE_HC | 16 | IgE_HC | 17 | 65 | K | SGGGVVQPGK | S | SLRLSBVASAFTFK | 2 |
| IgE_HC | 28 | IgE_HC | 362 | 13 | T | SLRLSBVASAFTFK | T | SPTITBLVVDLAPSK | 1 |
| IgE_HC | 30 | IgE_HC | 43 | 66 | K | SLRLSBVASAFTFK | K | RFAMHWVRQAPGK | 3 |
| IgE_HC | 43 | IgE_LC | 190 | 3 | K | RFAMHWVRQAPGKGLE | K | ADYEK | 1 |
| IgE_HC | 52 | IgE_LC | 151 | 3 | S | WVAVISDNGLRED | K | AKVQWKVDNALQSGNSQE | 2 |
| IgE_HC | 52 | IgE_LC | 161 | 6 | S | GLEWVAVISDNGLRED | S | AKVQWKVDNALQSGNSQE | 1 |
| IgE_HC | 65 | IgE_HC | 71 | 12 | K | DYEDSVK | S | GRFTISRDNSK | 1 |

| | | | | | | | | | |
|---|---|---|---|---|---|---|---|---|---|
| IgE_HC | 75 | IgE_LC | 111 | 2 | S | NSKDTLYLQmNSLRPED | T | RTVAAPSVFIFPPSDE | 1 |
| IgE_HC | 80 | IgE_LC | 147 | 5 | Y | NSKDTLYLQMNSLRPE | K | AKVQWKVDNALQSGNSQE | 1 |
| IgE_HC | 85 | IgE_LC | 188 | 3 | S | NSKDTLYLQMNSLRPED | Y | STYSLSSTLTLSKADYE | 1 |
| IgE_LC | 88 | IgE_LC | 95 | 59 | S | DFAVYSBQHYSD | S | SPPYTFGQGTK | 1 |
| IgE_LC | 123 | IgE_LC | 128 | 27 | S | RTVAAPSVFIFPPSDE | K | QLKSGTASVVBLLNNFYPRE | 1 |
| IgE_LC | 123 | IgE_LC | 133 | 17 | S | RTVAAPSVFIFPPSDE | S | QLKSGTASVVBLLNNFYPRE | 1 |
| IgE_LC | 123 | IgE_LC | 142 | 38 | S | RTVAAPSVFIFPPSDE | Y | QLKSGTASVVBLLNNFYPRE | 2 |
| IgE_LC | 128 | IgE_LC | 131 | 13 | K | RTVAAPSVFIFPPSDEQLK | T | SGTASVVBLLNNFYPREAK | 1 |
| IgE_LC | 133 | IgE_HC | 464 | 3 | S | SGTASVVBLLNNFYPREAK | K | KRTLABLIQNFmPE | 1 |
| IgE_LC | 151 | IgE_LC | 164 | 28 | K | VQWKVDNALQSGNSQE | S | SVTEQDSK | 2 |
| IgE_LC | 161 | IgE_HC | 512 | 7 | S | VDNALQSGNSQE | S | TKGSGFFVFSRLEVTRAEWE | 1 |
| IgE_LC | 164 | IgE_LC | 174 | 13 | S | VDNALQSGNSQESVTE | T | QDSKDSTYSLSSTLTLSK | 1 |
| IgE_LC | 164 | IgE_LC | 185 | 20 | S | VDNALQSGNSQESVTE | K | QDSKDSTYSLSSTLTLSK | 3 |
| IgE_LC | 164 | IgE_HC | 234 | 12 | S | SVTEQDSK | T | NKTFSVBSRDFTPPTVK | 1 |
| IgE_LC | 166 | IgE_LC | 185 | 77 | T | VDNALQSGNSQESVTE | K | QDSKDSTYSLSSTLTLSK | 3 |
| IgE_LC | 166 | IgE_HC | 234 | 14 | T | SVTEQDSK | T | NKTFSVBSRDFTPPTVK | 1 |
| IgE_LC | 205 | IgE_HC | 506 | 5 | S | HKVYABEVTHQGLSSPVTK | S | GSGFFVFSRLE | 1 |
| IgE_HC | 307 | IgE_HC | 315 | 31 | K | LTLSQKHWLSD | Y | RTYTBQVTYQGHTFE | 1 |
| IgE_HC | 307 | IgE_HC | 320 | 50 | K | LTLSQKHWLSD | T | RTYTBQVTYQGHTFEDSTK | 1 |
| IgE_HC | 552 | IgE_HC | 552 | 54 | K | AASPSQTVQRAVSVNPGK} | K | AASPSQTVQRAVSVNPGK} | 3 |
| **IgE-FcεRIα-Ara h 2_112** | | | | | | | | | |
| IgE_HC | 16 | IgE_LC | 182 | 4 | K | ESGGGVVQPGK | T | DSTYSLSSTLTLSKADYE | 1 |
| IgE_HC | 52 | IgE_HC | 69 | 4 | S | WVAVISDNGLREDYE | T | SVKGRFTISRDNSKD | 1 |
| IgE_HC | 52 | IgE_LC | 158 | 11 | S | GLEWVAVISDNGLRED | S | AKVQWKVDNALQSGNSQE | 3 |
| IgE_HC | 71 | IgE_HC | 78 | 40 | S | GRFTISRDNSK | T | DTLYLQMNSLRPE | 1 |
| IgE_HC | 80 | IgE_LC | 151 | 16 | Y | NSKDTLYLQMNSLRPE | K | AKVQWKVDNALQSGNSQE | 2 |
| IgE_HC | 85 | IgE_HC | 300 | 10 | S | DTLYLQmNSLRPE | S | LASTQSELTLSQK | 1 |
| IgE_LC | 88 | IgE_LC | 95 | 51 | S | DFAVYSBQHYSD | S | SPPYTFGQGTK | 3 |
| IgE_LC | 111 | IgE_LC | 131 | 56 | T | RTVAAPSVFIFPPSDE | T | QLKSGTASVVBLLNNFYPRE | 1 |
| IgE_LC | 123 | IgE_LC | 133 | 38 | S | RTVAAPSVFIFPPSDE | S | QLKSGTASVVBLLNNFYPRE | 3 |

| | | | | | | | | | |
|---|---|---|---|---|---|---|---|---|---|
| IgE_LC | 123 | IgE_LC | 142 | 11 | S | RTVAAPSVFIFPPSDE | Y | QLKSGTASVVBLLNNFYPRE | 2 |
| IgE_LC | 133 | IgE_HC | 401 | 3 | S | SGTASVVBLLNNFYPRE | T | EKQRNGTLTVTSTLPVGTRD | 1 |
| IgE_LC | 142 | IgE_LC | 151 | 72 | Y | SGTASVVBLLNNFYPRE | K | AKVQWK | 1 |
| IgE_LC | 151 | IgE_LC | 166 | 8 | K | VQWKVDNALQSGNSQE | T | SVTEQDSK | 2 |
| IgE_LC | 158 | IgE_LC | 185 | 26 | S | VDNALQSGNSQESVTE | K | QDSKDSTYSLSSTLTLSK | 1 |
| IgE_LC | 161 | IgE_HC | 466 | 8 | S | AKVQWKVDNALQSGNSQE | T | RTLABLIQNFMPE | 1 |
| IgE_LC | 164 | IgE_LC | 182 | 62 | S | VDNALQSGNSQESVTE | T | QDSKDSTYSLSSTLTLSK | 1 |
| IgE_LC | 164 | IgE_LC | 185 | 39 | S | VDNALQSGNSQESVTE | K | QDSKDSTYSLSSTLTLSK | 1 |
| IgE_LC | 164 | IgE_HC | 234 | 20 | S | SVTEQDSK | T | NKTFSVBSRDFTPPTVK | 1 |
| IgE_LC | 166 | IgE_LC | 185 | 25 | T | VDNALQSGNSQESVTE | K | QDSKDSTYSLSSTLTLSK | 3 |
| IgE_LC | 185 | IgE_LC | 192 | 48 | K | DSTYSLSSTLTLSKAD | K | YEKHK | 1 |
| IgE_LC | 190 | IgE_HC | 506 | 8 | K | YEKHK | S | TKGSGFFVFSRLEVTRAEWE | 1 |
| IgE_HC | 552 | IgE_HC | 552 | 57 | K | AASPSQTVQRAVSVNPGK} | K | AASPSQTVQRAVSVNPGK} | 2 |
| **IgE-FcεRIα-Ara h 2_221** | | | | | | | | | |
| IgE_HC | 7 | IgE_LC | 158 | 22 | S | SGGGVVQPGK | S | NALQSGNSQESVTE | 1 |
| IgE_HC | 16 | IgE_HC | 17 | 81 | K | ESGGGVVQPGK | S | SLRLSBVASAFTFK | 1 |
| IgE_HC | 16 | IgE_HC | 21 | 32 | K | ESGGGVVQPGK | S | SLRLSBVASAFTFK | 1 |
| IgE_HC | 16 | IgE_HC | 43 | 12 | K | SGGGVVQPGK | K | RFAmHWVRQAPGKGLE | 1 |
| IgE_HC | 30 | IgE_HC | 43 | 46 | K | SLRLSBVASAFTFK | K | RFAMHWVRQAPGK | 2 |
| IgE_HC | 52 | IgE_LC | 158 | 24 | S | GLEWVAVISDNGLRED | S | AKVQWKVDNALQSGNSQE | 2 |
| IgE_HC | 69 | IgE_HC | 80 | 22 | T | GRFTISRD | Y | NSKDTLYLQmNSLRPE | 1 |
| IgE_HC | 75 | IgE_LC | 158 | 3 | S | NSKDTLYLQMNSLRPE | S | AKVQWKVDNALQSGNSQE | 1 |
| IgE_HC | 85 | IgE_LC | 151 | 6 | S | NSKDTLYLQMNSLRPE | K | AKVQWKVDNALQSGNSQE | 1 |
| IgE_LC | 123 | IgE_LC | 128 | 25 | S | RTVAAPSVFIFPPSDE | K | QLKSGTASVVBLLNNFYPRE | 2 |
| IgE_LC | 123 | IgE_LC | 133 | 24 | S | RTVAAPSVFIFPPSDE | S | QLKSGTASVVBLLNNFYPRE | 1 |
| IgE_LC | 123 | IgE_LC | 142 | 16 | S | RTVAAPSVFIFPPSDE | Y | QLKSGTASVVBLLNNFYPRE | 3 |
| IgE_LC | 128 | IgE_LC | 131 | 16 | K | RTVAAPSVFIFPPSDEQLK | T | SGTASVVBLLNNFYPREAK | 1 |
| IgE_LC | 142 | IgE_LC | 151 | 20 | Y | SGTASVVBLLNNFYPRE | K | AKVQWK | 1 |
| IgE_LC | 142 | IgE_HC | 234 | 35 | Y | SGTASVVBLLNNFYPRE | T | WVDNKTFSVBSRDFTPPTVK | 1 |
| IgE_LC | 142 | IgE_HC | 311 | 13 | Y | SGTASVVBLLNNFYPRE | S | LASTQSELTLSQKHWLSD | 1 |

| | | | | | | | | | |
|---|---|---|---|---|---|---|---|---|---|
| IgE_LC | 151 | IgE_LC | 164 | 15 | K | VQWKVDNALQSGNSQE | S | SVTEQDSK | 2 |
| IgE_LC | 151 | IgE_LC | 166 | 79 | K | VQWKVDNALQSGNSQE | T | SVTEQDSK | 1 |
| IgE_LC | 158 | IgE_LC | 185 | 39 | S | VDNALQSGNSQESVTE | K | QDSKDSTYSLSSTLTLSK | 1 |
| IgE_LC | 164 | IgE_HC | 234 | 11 | S | SVTEQDSK | T | NKTFSVBSRDFTPPTVK | 1 |
| IgE_LC | 166 | IgE_LC | 175 | 79 | T | VDNALQSGNSQESVTE | Y | QDSKDSTYSLSSTLTLSK | 1 |
| IgE_LC | 166 | IgE_LC | 185 | 27 | T | VDNALQSGNSQESVTE | K | QDSKDSTYSLSSTLTLSK | 2 |
| IgE_LC | 166 | IgE_HC | 372 | 4 | T | NALQSGNSQESVTE | K | SPTITBLVVDLAPSK | 1 |
| IgE_LC | 170 | IgE_HC | 314 | 13 | S | SKDSTYSLSSTLTLSK | T | RTYTBQVTYQGHTFE | 1 |
| IgE_LC | 188 | IgE_HC | 496 | 8 | Y | YEKHKVYABE | S | VQLPDARHSTTQPRK | 1 |
| IgE_LC | 192 | IgE_LC | 209 | 27 | K | HKVYABE | K | VTHQGLSSPVTKSFNRGEB} | 2 |
| IgE_LC | 192 | IgE_LC | 210 | 38 | K | HKVYABEVTHQGLSSPVTK | S | SFNRGEB} | 1 |
| IgE_HC | 480 | IgE_HC | 503 | 10 | S | DISVQWLHNEVQLPD | T | TKGSGFFVFSRLEVTRAE | 2 |
| IgE_HC | 552 | IgE_HC | 552 | 67 | K | AASPSQTVQRAVSVNPGK} | K | AASPSQTVQRAVSVNPGK} | 3 |

**DETAILED METHODS:**

**Modeling the unbound bent / extended IgE and FcεRIα-bound bent IgE structures**

The IgE-Fc scaffolds (Cε1-Cε4) constructed previously[1] and referenced from PDB: 6EYO were used as templates to model bent and extended IgE, respectively. The Cκ was used as light chain constant domain. Sequences of Fv regions were substituted with fragments derived from published sequences (PA12C07) known for their high allergenicity to peanut allergens, as identified by Croote et. al[2]. Structures of these Fv regions, particularly CDR-H3, were modelled using ROSIE[3]. Full-length IgE models were reconstituted using MODELLER v10.1[4]. The FcεRIα-bound bent IgE complex was generated by superimposing FcεRIα at the FcεRIα-binding site located at the Cε3-Cε3 domains using PDB: 2Y7Q as template.

Glycans were added using CHARMM-GUI[5] (with Charmm36m forcefield) at six asparagine sites[6] (N144, N172, N222, N269, N375, and N398) on heavy Cε domains for the bent IgE and at five asparagines (excluding N375 due to mutation N375Q, referenced from PDB: 6EYO) for the extended IgE. These glycosylated models were solvated, minimized, and undergone production simulations for various replicates (Table 1). The glycosylated models were solvated with TIP3P water and neutralized with $K^+/Cl^-$ at 0.15 M concentration. Energy minimization (5000 steps using steepest descent) followed by a 50 ns equilibration with positional restraints were performed (400 kJ/mol nm² and 40 kJ/mol nm² to backbone and side chains, respectively). System temperature was maintained at 303.15 K using a Nosé-Hoover thermostat with a 1 ps time constant.

Production simulations were performed in replicates (Table S4) with different velocities. An isotropic Parrinello-Rahman barostat with a 5 ps time constant was employed to stabilize system pressure. Non-bonded and short-range electrostatic interactions were truncated at 0.9 nm through the potential shift Verlet cut-off scheme, while long-range electrostatic interactions were determined using the PME.

**Table S4:** Timescale and replicates of the IgE dynamics simulations

| Model | Abbreviations | Time scale × replicates |
|---|---|---|
| Unbound bent IgE | Ubent | 500 ns × 3 |
| Unbound extended IgE | Uextd | 500 ns × 3 |
| Ara h 2-bound IgE-Fab1 (region-I, interface-1) | *(refer Fig.S1-S4)* | 500 ns × 3 × 3 models |
| Ara h 2-bound IgE-Fab1 (region-II, interface-2) | *(refer Fig.S1-S4)* | 500 ns × 3 × 3 models |
| Ara h 2-bound bent IgE (region-III) | bent-Arah2 | 500 ns × 3 |
| [Ara h 2]$_2$-bound bent IgE (region-III) | *(refer Fig.S6)* | 500 ns × 3 |
| FcεRIα-bound bent IgE | FcR-bent | 500 ns × 3 |
| Arah2/FcεRIα-bound bent IgE | FcR-bent-Arah2 | 500 ns × 3 |

**Statistical analysis across different state ensembles**

For each of the 19 structural observables, *gap* was used to express the separation across the ensembles.

$$gap = \frac{\min(\left|mean_{target} - mean_{reference1}\right|, \left|mean_{target} - mean_{reference2}\right|)}{SD_{pooled}}$$

Where $mean$ was calculated based on 3 trajectory means per state and $SD_{pooled}$ was calculated within state trajectory-to-trajectory standard deviation. The Hedges' g for pairwise effect sizes and a multivariate PERMANOVA pseudo-F were then computed on the full 19-dimensional distances. Significance was assessed by using exact permutation of trajectory labels, i.e. 9! / (3!3!3!) = 1680 distinct relabeling. For two-group comparisons, all 10 distinct relabeling were used. Baseline *p-value* (at least one as extreme as observed) was calculated, ~0.0006 for three-group and 0.1 for any pairwise testing. Multiple testing across 19 observables was controlled by Benjamini-Hochberg false-discovery-rate (FDR) correction.

**Experimental preparation of IgE, Ara h 2, and FcεRIα**

IgE was produced and purified (by GenScript USA Inc., with Lot Number: U138NHK250-4/P9IB001) using published sequences of VH and VL (PA12C07), as identified by Croote et. al [2]. Constant domains included light chain Cκ and heavy chain Cε of human IgE.

Heavy chain:

QVQLEESGGGVVQPGKSLRLSCVASAFTFKRFAMHWVRQAPGKGLEWVAVISDNGLREDYEDSVKGRFTISRDNS

KDTLYLQMNSLRPEDTAIYYCAKVLDYSEYSLYFGMDVWGQGTTVLVSSASTQSPSVFPLTRCCKNIPSNATSVT

LGCLATGYFPEPVMVTWDTGSLNGTTMTLPATTLTLSGHYATISLLTVSGAWAKQMFTCRVAHTPSSTDWVDNKT

FSVCSRDFTPPTVKILQSSCDGGGHFPPTIQLLCLVSGYTPGTINITWLEDGQVMDVDLSTASTTQEGELASTQS

ELTLSQKHWLSDRTYTCQVTYQGHTFEDSTKKCADSNPRGVSAYLSRPSPFDLFIRKSPTITCLVVDLAPSKGTV

NLTWSRASGKPVNHSTRKEEKQRNGTLTVTSTLPVGTRDWIEGETYQCRVTHPHLPRALMRSTTKTSGPRAAPEV

YAFATPEWPGSRDKRTLACLIQNFMPEDISVQWLHNEVQLPDARHSTTQPRKTKGSGFFVFSRLEVTRAEWEQKD

EFICRAVHEAASPSQTVQRAVSVNPGK

Light chain:

EIVLTQSPAILSLSPGDRATLSCRTSQTVNSNFLAWYQQKPGQAPRLLIYGASRRAIDIPDRFTGSGSGTEFTLT

IARLEPEDFAVYSCQHYSDSPPYTFGQGTKLEIKRTVAAPSVFIFPPSDEQLKSGTASVVCLLNNFYPREAKVQW

KVDNALQSGNSQESVTEQDSKDSTYSLSSTLTLSKADYEKHKVYACEVTHQGLSSPVTKSFNRGEC

To note, there are a few differences in sequences of the purchased Ara h 2 and FcεRIα from those used in the computational modeling; however, the epitope regions on Ara h 2 and the IgE-binding sites on FcεRIα are reserved in both the sequences, as below:

```
Matrix: EBLOSUM62
 Gap penalty: 2.0
 Extend penalty: 2.0
 Score: 628.0
 Sequence 1 length:121
 Sequence 2 length:151
 Alignment length: 151
 Identity:      119/151 (78.81%)
 Similarity:    120/151 (79.47%)
 Gaps:           30/151 (19.87%)
                                    Region-I    Region-II
comp model: ---------RRCQSQLERANLRPCEQHLMQKIQRDEDSYERDPYSPSQ------------
                     |||||||||||||||||||||||||||||| ||||||||
purchased : RQQWELQGDRRCQSQLERANLRPCEQHLMQKIQRDEDSYGRDPYSPSQDPYSPSQDPDRR

                                                         Region-III
comp model: DPYSPSPYDRRGAGSSQHQERCCNELNEFENNQRCMCEALQQIMENQSDRLQGRQQEQQF
            ||||||||||||||||||||||||||||||||||||||||||||||||||||||||||||
purchased : DPYSPSPYDRRGAGSSQHQERCCNELNEFENNQRCMCEALQQIMENQSDRLQGRQQEQQF


comp model: KRELRNLPQQCGLRAPQRCDLD---------
            |||||||||||||||||||||
purchased : KRELRNLPQQCGLRAPQRCDLEVESGGRDRY
```

```
Matrix: EBLOSUM62
Gap penalty: 2.0
Extend penalty: 2.0
Score: 913.0
Sequence 1 length:170
Sequence 2 length:180
Alignment length: 180
Identity:      166/180 (92.22%)
Similarity:    166/180 (92.22%)
Gaps:           10/180 (5.56%)

comp model: ---KPKVSLNPPWNRIFKGENVTLTCNGNNFFEVSSTKWFHNGSLSEETNSSLNIVNAKF
               |||||||||||||||||||||||||||||||||||||||||||||||||||||||||
purchased : VPQKPKVSLNPPWNRIFKGENVTLTCNGNNFFEVSSTKWFHNGSLSEETNSSLNIVNAKF


comp model: EDSGEYKCQHQQVAESEPVYLEVFSDWLLLQASAEVVMEGQPLFLRCHGWRNWDVYKVIY
            ||||||||||||| ||||||||||||||||||||||||||||||||||||||||||||||
purchased : EDSGEYKCQHQQVNESEPVYLEVFSDWLLLQASAEVVMEGQPLFLRCHGWRNWDVYKVIY


comp model: YKDGEALKYWYENHAISITNAAAEDSGTYYCTGKVWQLDYESEPLNITVIKAP-------
            |||||||||||||| ||||||  ||||||||||||||||||||||||||||||
purchased : YKDGEALKYWYENHNISITNATVEDSGTYYCTGKVWQLDYESEPLNITVIKAPREKYWLQ
```

Sequence differences of Ara h 2 (above) and FcεRIα (below) between the computational model and the purchased. However, the epitope regions on Ara h 2 (highlighted in boxes) and the IgE-binding residues[7] on FcεRIα (shaded, referenced from PDB: 2Y7Q) are reserved in both the sequences.

**Protein Preparation and Buffer Exchange.**

Recombinant Ara h 2, human IgE, and FcεRIα were buffer exchanged into cross-linking buffer (50 mM HEPES, 100 mM NaCl, pH 7.4) using Amicon Ultra 0.5 mL centrifugal filters with a 3 kDa molecular weight cutoff (Merck Millipore). Protein concentrations were determined by measuring absorbance at 280 nm before and after buffer exchange to calculate protein loss. The final adjusted concentrations used for cross-linking were 3.77 mg/mL for IgE, 0.25 mg/mL for Ara h 2, and 1.84 mg/mL for FcεRIα.

**Sodium dodecyl sulfate-polyacrylamide gel electrophoresis (SDS-PAGE).**

A 10 µL aliquot from each cross-linking reaction was mixed with 4X LDS sample buffer, heated at 95°C for 10 minutes, and resolved on a 4-12% Bis-Tris polyacrylamide gel. The gel ran at 120 V for 30 minutes and subsequently stained using a Colloidal Blue Staining Kit (Invitrogen, LC6025) according to the manufacturer's instructions and then left to de-stain in milliQ water overnight.

**Cross-linking and sample preparation for mass spectrometry.**

Cross-linking was performed in 3 cross-link replicates with 4 different molar ratios: IgE:Ara h 2 (1:1), IgE:Ara h 2:FcεRIα (1:2:1 and 2:1:2), and IgE:FcεRIα (1:1). The mixtures (20 µg of IgE each) were incubated at 25°C for 15 minutes to allow complex formation. Cross-linking was initiated by adding 0.5 µL of DSSO from a 200 mM stock in dimethyl sulfoxide (DMSO), resulting in a final concentration of 2 mM DSSO and 1% DMSO. Proteins were incubated for 60 minutes at 25°C with shaking at 600 rpm. The reaction was quenched by adding 5 µL of 1 M Tris-HCl (pH 8.0) and incubating for a further 15 minutes. Negative controls for each complex mixture were prepared identically but without the addition of DSSO.

The cross-linked sample was then denatured with 50 µL of 8 M urea in 100 mM triethylammonium bicarbonate (TEAB), pH 8.5. Proteins were reduced by 10 mM Tris(2-carboxyethyl) phosphine (TCEP) for 20 minutes and alkylated with 55 mM chloroacetamide (CAA) in the dark for 30 minutes. The sample was then diluted with 100 mM TEAB to a final urea concentration below 2 M. A dual protease digestion was performed: first with LysC (1:25, w/w) for 4 hours at 25°C, followed by an overnight digestion with GluC (1:25, w/w) overnight at 25°C. The digestion was terminated by acidification to 1% trifluoroacetic acid (TFA). The resulting peptides were desalted using C18 StageTips (two C18 disk stacks per tip).

The StageTips were activated with 100 µL of 100% acetonitrile (ACN) and equilibrated twice with 100 µL of 0.1% formic acid (FA). Acidified peptide samples were loaded onto the tips, washed twice with 0.1% FA, and eluted with 100 µL of 65% ACN/0.1% FA. The eluted peptides were concentrated to dryness in a vacuum concentrator. The dried peptides were reconstituted in A*buffer to a final concentration of 0.5 µg/µL for LC-MS/MS.

**Cross-linking mass spectrometry.**

Peptide samples (2 µL per injection) were analyzed by liquid chromatography-tandem mass spectrometry (LC-MS/MS) on an Orbitrap Lumos mass spectrometer coupled to an EASY-nLC 1200 system (Thermo Fisher Scientific). Peptides were separated on a 75 µm x 50 cm C18 column using a 60-minute linear gradient from 3% to 30% mobile phase B (80% ACN, 0.1% formic acid) at a flow rate of 300 nL/min. MS2-MS2 fragmentation was used, with MS1 acquisition performed in Orbitrap with 60,000 resolution for a scan range of 400-1600 m/z with AGC target of 400,000 and maximum injection time of 118 ms in profile mode. MS2 acquisition was performed in Orbitrap with 30,000 resolution with CID activation with collision energy of 30 %. AGC target of 50,000 and maximum injection time of 100 ms was set. MS2 acquisition was performed in Orbitrap with HCD activation with collision energy of 35 % and 30,000 resolution, AGC target of 50,000, and maximum injection time of 100 ms.

**Analysis of cross-linking mass spectrometry data.**

The raw data were processed using the MeroX software[8] (version 2.0.1.4) to identify cross-linked peptides. Data were searched against a database containing sequences of Ara h 2, IgE heavy and light chains, and FcεRIα. A calibration search with precursor mass tolerance of 10.0 ppm and product mass tolerance of 20.0 ppm was performed before the cross-link search. Search was performed with DSSO cross-link on K, S, T, Y amino acids with a maximum of 3 missed cleavages, LysC/GluC proteases, and fixed modification for carbamidomethyl (C) and variable modifications for oxidation (M), deamidation (N, Q). The false discovery rate (FDR) cut-off was set to 1.0%.

**Hydrogen-Deuterium Exchange Mass Spectrometry (HDX-MS)**

HDX-MS were performed in four setups: apo Ara h 2 (90pmol), apo IgE (90pmol), Ara h 2-IgE complex (180pmol: 90pmol), and FcεRIα-IgE-Ara h 2 complex (90pmol: 90pmol:180pmol). For complex formation, Ara h 2 and IgE or FcεRIα-IgE-Ara h 2 were pre-incubated at room temperature for 30 min prior to deuterium labeling.

Deuterium exchange buffer was prepared by lyophilizing the protein sample buffer 50 mM HEPES pH7.4, 100 mM NaCl to remove $H_2O$ and reconstituting with 99.9% $D_2O$ (Waters, Milford, MA). Exchange reactions were initiated by diluting the protein or protein complex into 10-fold excess of deuterium buffer, yielding a final $D_2O$ concentration of 90%. Labeling was conducted at three timepoints: 1 min, 10 min, and 100 min.

HDX reactions were quenched by lowering the pH to 2.5, achieving a final concentration of 1.5 M guanidine hydrochloride (GnHCl), in the presence of tris(2-carboxyethyl) phosphine (TCEP) to reduce disulfide bonds. Quenching was performed for 1 min on ice to minimize back-exchange. All labeling experiments were performed in triplicate, and reported deuterium uptake values represent the mean of replicates without correction for back-exchanges.

Quenched samples were analyzed using a nano-UPLC HDX system (Waters, Milford, MA, USA). Online digestion was performed on a Waters Enzymate BEH pepsin column (2.1 × 30 mm) with 0.1% formic acid in water at 100 µL/min. Peptic fragments were trapped on a C18 VanGuard pre-column (2.1 × 5 mm, 1.7 µm) and separated on an ACQUITY UPLC BEH C18 column (1.0 × 100 mm, 1.7 µm) using an 8–40% acetonitrile gradient in 0.1% formic acid at 40 µL/min. Peptides were ionized by electrospray and analyzed on a SYNAPT G2-Si mass spectrometer (Waters, Milford, MA, USA) in MSE mode. Continuous calibration was achieved by co-infusion of [Glu1]-fibrinopeptide B (100 fmol/µL) at 10 µL/min.

ProteinLynx Global Server (PLGS) v3.0 was used to identify the sequences from mass spectra data based on undeuterated protein samples. Separate protein sequence database of each protein sequence was used for peptide identification. Following search parameters were used in PLGS search:(i) no specific protease and (ii) variable N-linked glycosylation modification. Additional cutoff filters applied included (i) minimum intensity = 1000, (ii) minimum products per amino acids = 0.1, and (iii) a precursor ion mass tolerance of <10 ppm in DynamX v.3.0 (Waters, Milford, MA). Peptides independently identified under the specified condition and present in at least in two out of three undeuterated sample replicates were retained for HDX-MS analysis. Uptake per peptide was calculated as the difference between the centroid mass of labeled peptides (1, 10, and 100 min) and the unlabeled state. Deuteros 2.0 software was used to perform statistical analysis on the differences in deuterium exchange among the identified peptides. Comparative analyses between free and bound states (Ara h 2-IgE and FcεRIα-IgE-Ara h 2) were expressed as difference plots across the protein sequence using

Woods plots and a confidence interval (CI) of 99% was used to identify peptides showing a significant difference.

**Table S5:** HDX-MS experimental summary data table

| | **Ara h2** | **IgE** | **Ara h2- IgE** | **IgE –FcϵRIα** | **Ara h2 – IgE – FcϵRIα** |
|---|---|---|---|---|---|
| HDX reaction details | 50mM HEPES, 100mM NaCl in 90% $D_2O$, pH 7.2, 37°C | 50mM HEPES, 100mM NaCl in 90% $D_2O$, pH 7.2, 37°C | 50mM HEPES, 100mM NaCl in 90% $D_2O$, pH 7.2, 37°C | 50mM HEPES, 100mM NaCl in 90% $D_2O$, pH 7.2, 37°C | 50mM HEPES, 100mM NaCl in 90% $D_2O$, pH 7.2, 37°C |
| HDX time points (mins) | 1, 10 and 100 | | | | |
| Quench buffer and conditions | Addition of 0.1% TFA, 1.5 M guanidine hydrochloride (GnHCl) , tris(2-carboxyethyl) phosphine (TCEP) and 1 min incubation on ice | | | | |
| HDX controls | Maximally labelled controls were not performed | | | | |
| Back-exchange correction | No done | | | | |
| No. of peptides | 8 | 16/12 | 16/12 | 12/11 | 10/12 |
| Sequence coverage (%) | 32.8 | 44.4/54.9 | 44.4/54.9 | 28.4/52.1 | 30.6/54.8 |
| Average peptide length / redundancy | 18.3/2.9 | 16.9/1.1<br>12.8/1.3 | 16.9/1.1<br>12.8/1.3 | 15.2/1.2<br>13.1/1.3 | 16.9/1.1<br>12.75/1.3 |
| Replicates | 3 (Technical) | 3 (Technical) | 3 (Technical) | 3 (Technical) | 3 (Technical) |
| Repeatability | 0.103 | 0.061/0.036 | 0.061/0.036 | 0.075/0.040 | 0.049/0.038 |
